# CDK8 and CDK19 act redundantly to control the CFTR pathway in the intestinal epithelium

**DOI:** 10.1101/2022.01.28.478171

**Authors:** Susana Prieto, Geronimo Dubra, Alain Camasses, Elisabeth Simboeck, Ana Bella Aznar, Christina Begon-Pescia, Nelly Pirot, François Gerbe, Lucie Angevin, Philippe Jay, Liliana Krasinska, Daniel Fisher

## Abstract

CDK8 and CDK19 form a conserved cyclin-dependent kinase subfamily that interacts with the essential transcription complex, Mediator, and also promotes transcription by phosphorylating the C-terminal domain (CTD) of RNA polymerase II. Cells lacking either CDK8 or CDK19 are viable and have limited transcriptional alterations, but whether the two kinases redundantly control cell differentiation is unknown. Here, we find that CDK8 is dispensable for RNA polII CTD phosphorylation, regulation of gene expression, normal intestinal homeostasis and efficient tumourigenesis in mice. Furthermore, CDK8 is largely redundant with CDK19 in the control of gene expression. Yet, while their combined deletion in intestinal organoids reduces long-term proliferative capacity, it is not lethal and allows differentiation. Nevertheless, in double mutant organoids, the cystic fibrosis transmembrane conductance regulator (CFTR) pathway is transcriptionally and functionally downregulated, leading to mucus accumulation and increased secretion by goblet cells. This phenotype can be recapitulated by pharmacological inhibition of CDK8/19 kinase activity. Thus, the Mediator kinases are not essential for cell proliferation and differentiation, but they cooperate to regulate tissue-specific transcriptional programmes.

## Introduction

CDK8 was discovered as a kinase that binds cyclin C and, like CDK7-cyclin H and CDK9-cyclin T, can promote transcription by phosphorylating the C-terminal repeat domain (CTD) of RNA polymerase II (PoI II) (Rickert *et al*, 1999). CDK8 and cyclin C are exceptionally highly conserved in vertebrates, as illustrated by 97% and 98% amino acid identity over the whole sequence between *Xenopus* and human CDK8 and cyclin C, respectively. This unusual level of cross-species conservation implies critical functions in fundamental cellular processes. In complex with Med12 and Med13, CDK8-cyclin C forms the canonical cyclin-dependent kinase module (CKM) of the Mediator transcriptional co-regulator complex, a function that is conserved with the more divergent yeast homologues of CKM subunits (Jeronimo *et al*, 2016). The latter were revealed as suppressors of a CTD truncation, suggesting a transcription-repressive activity of the CKM (Liao *et al*, 1995). In vertebrates, a second member of the CDK8 subfamily, CDK19, almost identical in the kinase domain with CDK8, also binds cyclin C and interacts with Mediator, in a manner generally thought to be exclusive with CDK8 (Sato *et al*, 2004; Tsutsui *et al*, 2008; Knuesel *et al*, 2009).

Mediator is a large multi-subunit complex required for Pol II-dependent transcription in all eukaryotes (Malik & Roeder, 2010). Acute ablation of vertebrate Mediator is lethal for cells and results in a rapid downregulation of the entire transcriptome (El Khattabi *et al*, 2019). The tail subunits integrate enhancer-associated transcription factor activity into conformational changes of the head- and middle complex, which controls Pol II interactions with the basal transcriptional machinery at promoters as well as phosphorylation of the CTD (Malik & Roeder, 2010). Biochemical analysis in yeast provided evidence that the CKM negatively regulates Mediator. *In vitro* experiments suggest that it hinders basal transcription by sterically blocking CTD-dependent recruitment of PolII to Mediator middle subunits (Elmlund *et al*, 2006; Tsai *et al*, 2013); while *in vivo* data show that the CKM binds to the same promoters as core Mediator but with low stoichiometry (Andrau *et al*, 2006). This appears to be due to negative regulation of Mediator binding to upstream enhancer sequences and release of the CKM module upon Mediator-Pol II interactions (Jeronimo *et al*, 2016).

In contrast to Mediator, the activity of the CKM is apparently non-essential in many cell types, as genes encoding CDK8, CDK19 and cyclin C are not required for survival and proliferation of most cell types in different organisms (Loncle *et al*, 2007; Kuchin *et al*, 1995; Li *et al*, 2014; Postlmayr *et al*, 2020). However, CDK8 is required for normal development. Germline ablation of *Cdk8* is lethal at the pre-implantation stage in mice (Westerling *et al*, 2007), while conditional deletion using a *Sox2* Cre driver is lethal around embryonic day 10.5 (Postlmayr *et al*, 2020). The difference in lethality stage between the two genotypes suggests that CDK8 might be essential in zygotes, prior to *Sox2* expression.

Deletions of other CKM subunits in mice have variable phenotypes. Cyclin C gene deletion is embryonic lethal at day 10.5 with severe growth defects, and its deletion in adults affects T-cell differentiation (Li *et al*, 2014), while deletion of *Med12* is lethal at late embryonic stages, preventing neural-tube closure, axis elongation and organ morphogenesis (Rocha *et al*, 2010). CDK19 deletion has not yet been reported. An essential requirement for CKM subunits in transcriptional regulation in animals cannot, however, be completely ruled out, since differences in the lethality stage of CKM subunit deletions could be due to differential maternal mRNA contributions.

Consistent with a repressive role for the CKM in transcription, we recently reported that inhibition of CDK8 and CDK19 in human and mouse pluripotent stem cells is associated with a global overactivation of enhancers and a stabilisation of the naiive state (Lynch *et al*, 2020, 8). Similarly, in acute myeloid leukaemia, CDK8/19 bind superenhancers and their chemical inhibition further activates enhancer activity (Pelish *et al*, 2015).

CDK8 has been attributed oncogenic functions in different cancers, including Wnt-dependent colorectal cancer, melanoma, breast and prostate cancer, acute myeloid leukaemia and B-cell leukaemia (Pelish *et al*, 2015; Firestein *et al*, 2008; Morris *et al*, 2008; Kapoor *et al*, 2010; McDermott *et al*, 2017; Nakamura *et al*, 2018; Menzl *et al*, 2019). Originally proposed to act in intestinal cancers by promoting Wnt transcription (Firestein *et al*, 2008), CDK8 is also involved in transcription dependent on Notch signalling (Li *et al*, 2014), NFκB (Chen *et al*, 2017), HIF1α (Galbraith *et al*, 2013), the serum-response (Donner *et al*, 2010), the interferon-_γ_ response (Bancerek *et al*, 2013; Steinparzer *et al*, 2019), p53 (Donner *et al*, 2007), superenhancers (Pelish *et al*, 2015), histone variant incorporation into chromatin (Kapoor *et al*, 2010), in pluripotency maintenance (Adler *et al*, 2012) and the senescence-associated tumour-promoting secretory phenotype (Porter *et al*, 2012). It also restrains NK-mediated cell toxicity and tumour surveillance (Hofmann *et al*, 2020). CDK8 is thought to act either by directly phosphorylating transcription factors such as Notch1 (Li *et al*, 2014) or STAT1 (Bancerek *et al*, 2013), or by being co-recruited to promoters via specific transcription factors, where it phosphorylates PolII CTD (Chen *et al*, 2017; Galbraith *et al*, 2013; Steinparzer *et al*, 2019). Yet genetic confirmation of requirements for CDK8 in cancers *in vivo* has been lacking. Conditional knockout of *Cdk8* in the intestinal epithelium did not hinder intestinal tumour development in *Apc* mutant mice; rather, it appeared to enhance tumourigenesis, and knockouts were reported to have lost the Polycomb group 2-mediated repressive histone mark, H3 lysine-27 trimethylation, thus upregulating oncogenic transcription (McCleland *et al*, 2015).

In contrast to CDK8, almost nothing is known about CDK19 roles in cancer, and whether it compensates for loss of CDK8 remains unknown. *In vitro* inhibition or knockdown experiments have suggested that CDK8 and CDK19 control different sets of target genes (Tsutsui *et al*, 2008, 11; Galbraith *et al*, 2013, 11; Poss *et al*, 2016). By genetic ablation in mice liver cells, we recently found that CDK8 and CDK19 are both required for hepatic carcinogenesis, and highlighted genetic interaction with p53 as critical for their roles in tumourigenesis (Bacevic *et al*, 2019).

A number of potent CDK8 pharmacological inhibitors have been developed (Pelish *et al*, 2015; Porter *et al*, 2012; Hofmann *et al*, 2020; Bergeron *et al*, 2016; Koehler *et al*, 2016; Dale *et al*, 2015; Schiemann *et al*, 2016), which are expected to also target CDK19. Anti-cancer activity of CDK8/19 inhibitors has been somewhat limited, and there may be only a small therapeutic window, due to systemic toxicity. However, debate about whether CDK8/19 inhibitor toxicity is on-target, *i.e*. due to inhibition of CDK8 and CDK19 (Clarke *et al*, 2016), or off-target, due to inhibition of other kinases (Chen *et al*, 2019), continues. Furthermore, at least some CDK8/19-mediated phenotypes appear to be kinase-independent (Steinparzer *et al*, 2019; Audetat *et al*, 2017; Menzl *et al*, 2019).

Thus, despite their established roles as regulators of Mediator, and considerable interest in their therapeutic targeting in cancer, we do not yet fully understand the redundant and specific roles of CDK8 and CDK19. We therefore used gene-targeting in mice to address these questions, and determine whether their combined deletion is lethal. We confirm that knockout of the *Cdk8* gene in the intestinal epithelium has little or no effect on cell proliferation or differentiation. Furthermore, double deletion of *Cdk8* and *Cdk19* is compatible with cell proliferation in intestinal organoids. However, the double knockout reveals redundant functions in long-term control of cell proliferation and gene expression programmes. We uncover an unexpected requirement for these kinases in control of the CFTR pathway, a key player in cystic fibrosis.

## Results and discussion

To evaluate the possible requirements for CDK8 for cell proliferation and survival in adult vertebrates, we designed and generated a conditional knockout allele of *Cdk8* in the mouse by Lox/Cre targeting exon 2 (Fig S1A, B). This removes the critical catalytic lysine-52 and results in a frameshift that truncates over 90% of the protein. A similar conditional *Cdk8* allele was independently generated (McCleland *et al*, 2015). We studied the requirement for CDK8 in the adult mouse intestine since this is one of the most highly proliferative tissues in adults. We crossed *Cdk8*^*lox/lox*^ mice to mice expressing a Tamoxifen-inducible Cre under the control of the Villin promoter (el Marjou *et al*, 2004), and verified efficient deletion of *Cdk8* in the intestinal epithelium by genotyping and Western blotting (Fig S1C, S2B). Mice lacking CDK8 were healthy and did not present any phenotypes in the intestine; there was no difference in the number of proliferating cells nor in cell cycle distribution as assessed by the number of BrdU positive cells after a two-hour pulse (Fig 1A, B, Fig S2A). The number of stem cells, goblet cells, tuft cells and Paneth cells was similar to wild-type mice, indicating that differentiation programmes were not affected (Fig 1B, Fig S2A). The deletion was maintained after 2 months, showing that there is no counter-selection for non-recombined intestinal crypts (Fig S2B). We performed RNA-sequencing from the intestinal epithelium of wild-type and knockout mice, but could identify only 4 genes (*Wdr72, Trim12a, Npr3, Trim30d*) with statistically significant expression alterations (all of which were upregulated), suggesting that CDK8 loss has only minor effects on gene expression in the intestine that are obscured by biological variability between animals.

**Fig. 1.**
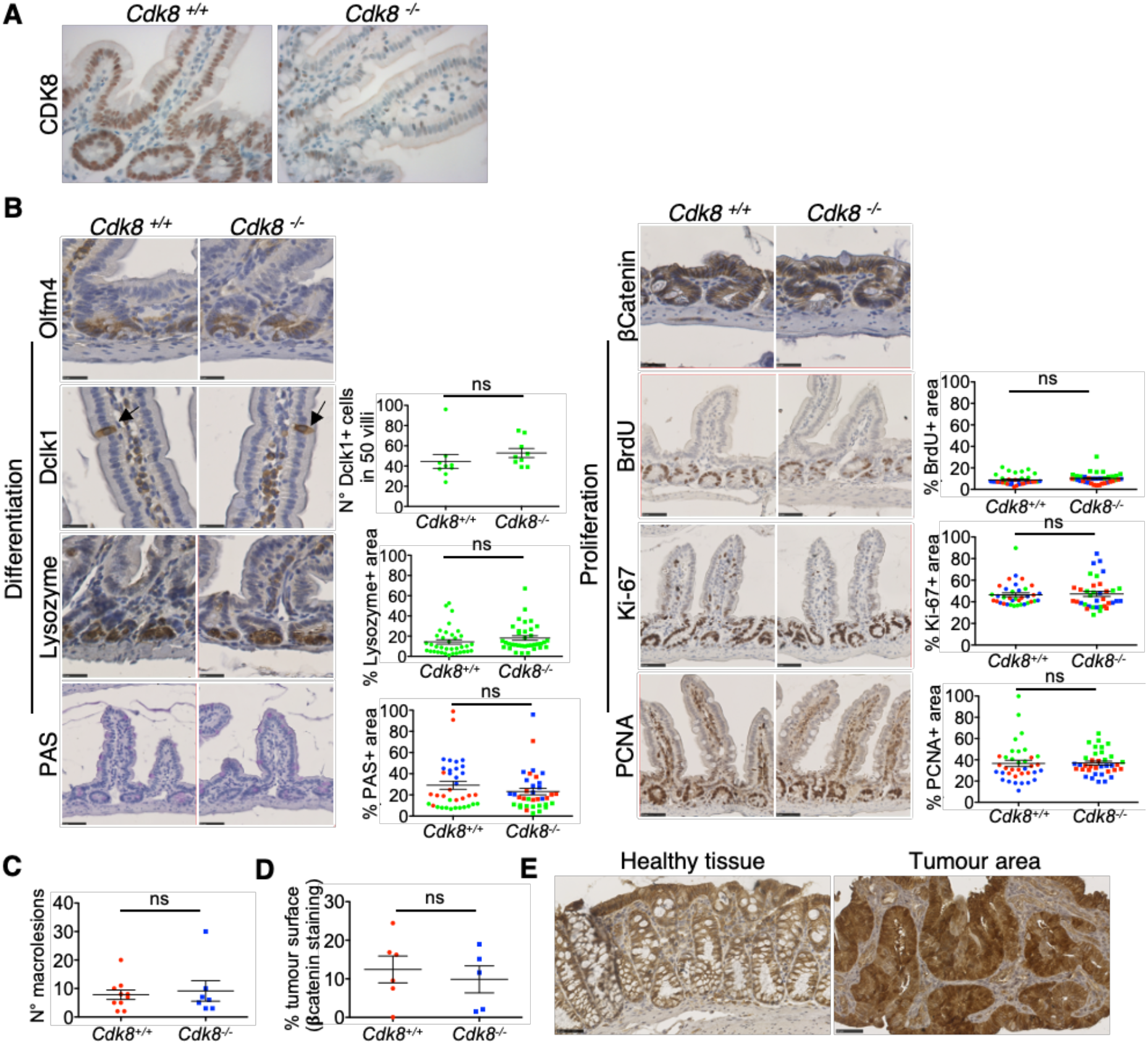
CDK8 knockout does not affect adult mouse intestine homeostasis nor chemically-induced carcinogenesis. **(A)** Immunohistochemical staining of CKD8 in mouse small intestines collected two months after tamoxifen treatment. CDK8^+/+^ depict mice with floxed Cdk8 alleles. (B) Analysis of cell differentiation (left) and proliferation (right) in the intestine after CDK8 deletion (as in A). Olfm4, Lysozyme, PAS and Dclk1 staining was used to reveal, respectively, stem, Paneth, goblet and tuft cells. β-Catenin staining allows detection of cancer cells (cytoplasmic vs nuclear localisation). Cell proliferation was assessed by PCNA, Ki-67 and BrdU (after 1h pulse) staining. Scatter plots represent the percentage of the area stained by each antibody (relative to the area occupied by hematoxylin). For Paneth cells, BrdU, PCNA and Ki67, only crypts were analysed. For goblet cells, crypts and villi were analysed. For Tuft cells quantification, Dclk1 positive cells were counted in 50 villi. Colour code depicts small intestine (green), proximal colon (blue), and distant colon (red). Mean ± SEM is shown. P-value of unpaired two-tailed t-test is indicated (ns, not significant; p > 0.05). Scale bars, 25μm (Olfm4, Lysozyme, Dclk1 and β-Catenin) and 50μm (PAS, BrdU, Ki-67 and PCNA). (C-E) Analysis of mouse colon after AOM/DSS treatment. (C) Quantification of the number of neoplastic lesions (n = 10 for Cdk8 ^+/+^, and n = 7 for Cdk8 ^-/-^ mice). P-value of umpaired t-test is indicated: ns, not significant (p > 0.05). Mean ± SD is shown. (D) Quantification of the percentage of the colon surface occupied by tumours. Intestine samples were stained for β-Catenin, and tumour regions with nuclear β-Catenin localisation were quantified using NDP.view software. Two-tailed p-value of unpaired t-test is indicated; ns, not significant (p > 0.05). Mean ± SD is shown. **(E)** Example of IHC with β-Catenin staining of tumour-free regions with membrane β-catenin localization *(left)* and tumour regions with nuclear β-catenin localisation *(right)*. Scale bars, 50μm.

CDK8, like CDK7, can phosphorylate PolII CTD (Rickert *et al*, 1999). Since cyclin C deletion in mice abolishes CDK8 activity yet does not affect PolII CTD phosphorylation (Li *et al*, 2014), while CTD S5 phosphorylation is normal in a non-proliferative tissue (liver) of *Cdk7* knockout mice (Ganuza *et al*, 2012), we asked whether CDK7 and CDK8 can compensate each other in CTD phosphorylation. We crossed single floxed *Cdk7* and *Cdk8* mice to generate *Cdk7*^*lox/lox*^; *Cdk8*^*lox/lox*^ mice, with Cre-expressed under control of a ubiquitous promoter (RPB1). While complete CDK8 loss occurred in both intestine and liver, CDK7 loss was incomplete in both tissues (Fig S3A, B). In the intestine this incomplete deletion was expected since CDK7 is required for cell proliferation; thus, rare non-recombined crypts repopulate the epithelium (Ganuza *et al*, 2012). The strong reduction of CDK7 combined with ablation of CDK8 only led to a slight reduction in phosphorylation of PolII CTD S5, while S2 and S7 phosphorylation were normal (Fig S3A, B). These results do not indicate a critical role for CDK8 in this phosphorylation *in vivo*, and show that a low level of CDK7 is sufficient for PolII CTD-phosphorylation.

CDK8 was described as an oncogene in colorectal cancer where it promotes beta-catenin-dependent transcription (Firestein *et al*, 2008; Morris *et al*, 2008), suggesting that its deletion should inhibit intestinal tumorigenesis. To test this, we used a colitis-associated chemical model of intestinal tumourigenesis. We treated adult control mice or mice lacking *Cdk8* with azoxymethane-dextran sodium sulphate (AOM-DSS) to chemically induce intestinal tumours, and sacrificed mice 2 months after the first DSS treatment (Fig S4A). We observed no effect of *Cdk8* deletion on colitis-induced weight loss (Fig S4B) and no counter-selection for *Cdk8* deletion in the tumours (Fig S4C, D). There was no difference in number or area of tumours between WT and knockout animals (Fig 1C-E). We next tested whether CDK8 loss affects acute activation of the Wnt pathway, by concomitant homozygous deletion of the *Adenomatous polyposis coli* (*Apc*) tumour suppressor gene. Both *Apc*^*-/-*^; *Cdk8*^*+/+*^and *Apc*^*-/-*^; *Cdk8*^*-/-*^mice showed rapid morbidity necessitating sacrifice 5 days after Tamoxifen treatment, and hyperplastic intestinal epithelium. Intestinal epithelium lacking both *Cdk8* and *Apc* had no difference in number of Ki-67-positive cells when compared to mutant *Apc* alone, indicating similar cell proliferation. (Fig S5A-D). Taken together, our data do not support a major role for CDK8 in intestinal tumourigenesis in the mouse.

The lack of striking phenotypes of *Cdk8* deletion in the adult intestine, along with the almost complete conservation of the kinase domain between CDK8 and CDK19 and across vertebrate species (Fig S6A) suggested that if CDK8 has essential functions, CDK19 might be able to compensate for its loss. We therefore sought to generate a conditional double knockout. Since double *Cdk8/Cdk19* knockout in adult mice might be lethal, we undertook a conditional deletion using intestinal organoids, which recapitulate many features of intestinal development and morphology while facilitating genetic manipulation *in vitro* (Clevers, 2016). We thus generated intestinal organoids from WT and *Cdk8*^*lox/lox*^; *Vill::Cre*^*ERT2/+*^ mice and disrupted *Cdk19* by CRISPR-Cas9-directed gene targeting (Fig S6B), using retroviral transduction of Cas9 and a synthetic guide RNA-expressing plasmid. *Cdk8* removal was efficient after 7 days of Tamoxifen treatment (Fig 2A, B). Cyclin C was lost specifically in double knockout organoids (Fig 2B) and correlated with a loss of STAT1 phosphorylated on S727, a previously described CDK8 substrate (Bancerek *et al*, 2013, 727), confirming redundancy of the two kinases. Growth appeared somewhat slower only in double knockout organoids (Fig 2C, D), with a corresponding larger fraction of non-proliferating cells, as demonstrated by loss of Ki-67 staining (Fig 2E, F). Consistent with slower growth, *Cdk8* deletion was counter-selected in *Cdk19* knockout organoids, as long-term culture resulted in reappearance of CDK8 in two out of three double knockout organoid populations, presumably due to expansion of a minor unrecombined population (Fig S7).

**Fig. 2.**
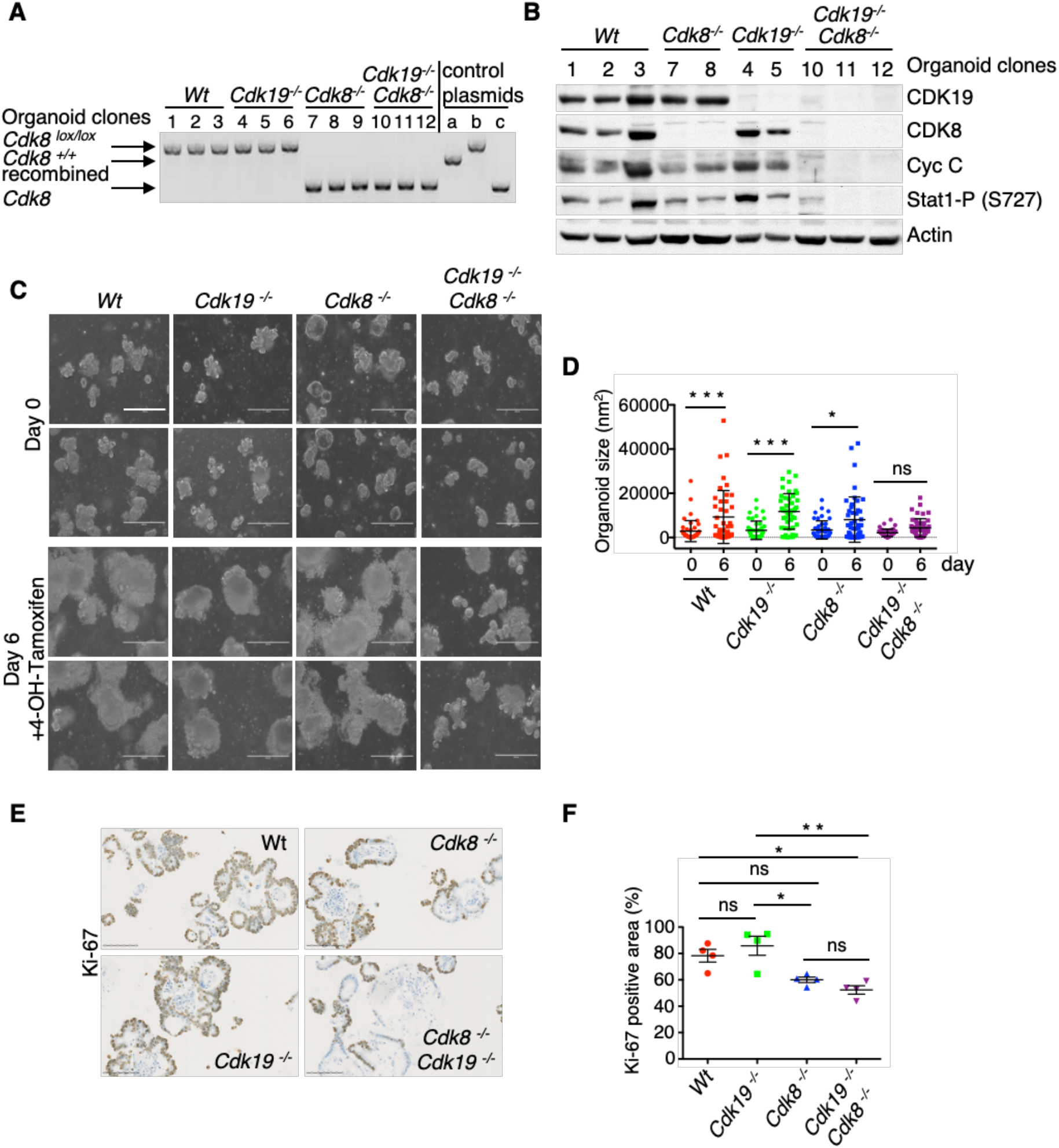
Double CDK8/CDK19 knockout intestinal organoids show decreased cell proliferation. **(A)** Genotyping confirms the loss of *Cdk8* exon 2 in *Cdk8* ^*-/-*^ and *Cdk8* ^*-/-*^*/Cdk19* ^*-/-*^ organoids after 7 days of OH-tamoxifen treatment. Control plasmids (a, b and c) are described in Fig S1C. **(B)** WB of organoid samples after 7 days of OH-tamoxifen treatment; β-actin was used as loading control. **(C)** Phase contrast images of organoids before and after 6 days of OH-tamoxifen treatment. Scale bars, 150μm. **(D)** Quantification of organoid size at day 0 and 6, as in B (mean + SD are shown). P-value, ordinary one-way ANOVA: (*) p ≤ 0.05, (***) p ≤ 0.001. **(E)** IHC staining of organoids (day 7 of OH-tamoxifen treatment) with Ki-67 antibody. Scale bars, 100μm. **(F)** Quantification of Ki-67 positive area (% of the total area of the organoids; mean + SD) in the four different genotypes presented in (E). Areas with positive Ki-67 signal were detected and quantified using QuPath and ImageJ programs. Adjusted p-values of ordinary one-way ANOVA followed by Tukey’s multiple comparison test are indicated: (***) p-value ≤ 0.001; (**) p-values ≤ 0.01; (*) p-values ≤ 0.05; ns, not significant (p > 0.05).

To determine effects of loss of the Mediator kinases on gene expression, we performed RNA-sequencing analysis of stable populations of single and double *Cdk8/Cdk19* knockout organoids. CDK8 loss had a stronger effect (716 genes upregulated, 575 downregulated) than CDK19 loss (158 up, 151 down), while double knockout organoids (1819 up, 1363 down; Fig 3A, B) revealed functional redundancy between CDK8 and CDK19 in regulating gene expression. However, most expression alterations were minor, with only 830 genes deregulated by a factor of two or more. This is consistent with previous studies, none of which have shown sweeping changes in the transcriptome upon downregulation or inhibition of CDK8 or CDK19, but rather, alteration of a limited number of specific gene sets, including super-enhancer associated genes (El Khattabi *et al*, 2019; Pelish *et al*, 2015; Galbraith *et al*, 2013; Steinparzer *et al*, 2019; Poss *et al*, 2016; Clarke *et al*, 2016). In terms of genes controlling cell proliferation, cyclin A (*Ccna2)*, cyclin B (*Ccnb2)* and cyclin E (*Ccne2*) were slightly downregulated in knockouts, but this is likely to be a consequence rather than a cause of reduced cell proliferation. CDK8 has previously been found to regulate the p53 and c-Myc pathways (Donner *et al*, 2007; Adler *et al*, 2012), but intestinal cells lacking both kinases showed only a slight (though significant) downregulation of c-Myc, while p53 was not affected. In contrast, cyclin G1, a positive mediator of p53 responses and RB functions with a role in cell cycle arrest (Zhao *et al*, 2003), and p21 (*Cdkn1A)*, a p53 target that inhibits cyclin-dependent kinases to provoke cell cycle arrest, were more strongly upregulated in the double knockout organoids.

**Fig. 3.**
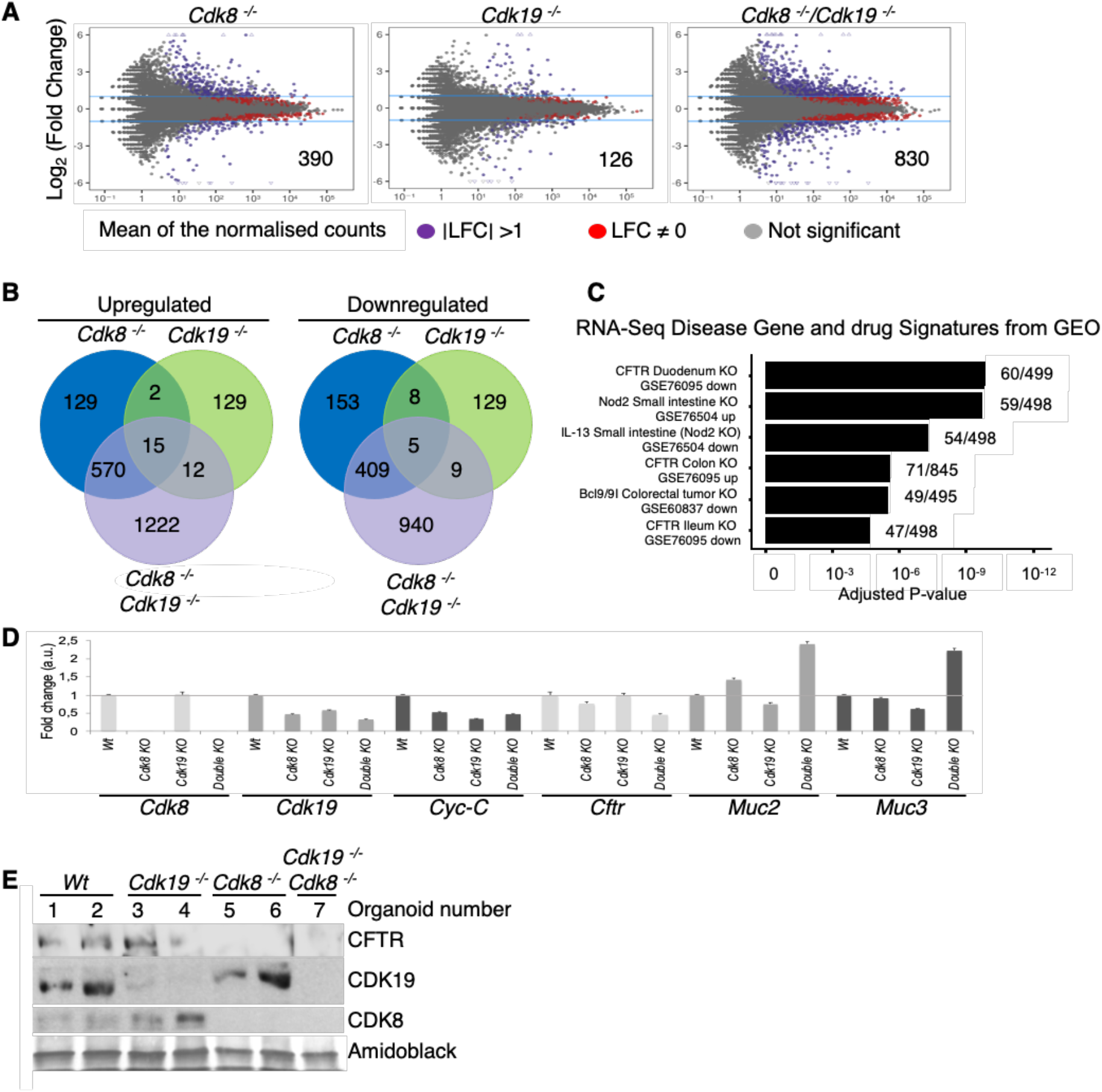
Functional redundancy between CDK8 and CDK19 in regulation of gene expression. **(A)** Dot plot analysis of differentially expressed genes (DEGs). Red dots: DEGs with p-value ≤ 0.05; purple dots: log_2_ fold change (LFC) >1 or <-1, p-value ≤ 0.05; grey dots: not significant, NS. Numbers inside plots indicate the number of genes deregulated more than 2-fold. **(B)** Venn diagrams indicating intersection of genes with altered expression in the indicated genotypes. **(C)** Gene set enrichment analysis (using Enrichr database) of highly deregulated genes in *Cdk8*^*-/-*^*/Cdk19*^*-/-*^ organoids. Manually curated signatures extracted from RNA-seq studies in GEO where gene expression was measured before and after drug treatment, gene perturbation or disease. **(D)** qRT-PCR analysis of indicated mRNA levels in Wt, *Cdk8*^*-/-*^, *Cdk19*^*-/-*^ and *Cdk8*^*-/-*^*/Cdk19* ^*-/-*^ organoids. **(E)** WB of indicated proteins extracted from Wt, *Cdk8* ^*-/-*^, *Cdk19* ^*-/-*^ and *Cdk8* ^*-/-*^*/Cdk19* ^*-/-*^ organoids. Amidoblack was used as loading control.

Pathway analysis in double knockout organoids unexpectedly revealed a significant alteration of genes also modulated in intestinal knockouts of the cystic fibrosis transmembrane conductance regulator, CFTR (Fig 3C). In double knockouts, expression of genes involved in mucus production, *Muc2, Muc3, Muc13, Nlrp6, Agr2, Gcnt4, Tff1*, were upregulated, while *Cftr* was reduced. We validated changes of selected genes by qRT-PCR (Fig 3C, D). The loss of *Cftr* mRNA was also reflected at the protein level, since CFTR protein was lost in double mutant organoids (Fig 3E).

Cystic fibrosis is a disease of mucosal epithelia which also affects the intestine, and is characterised by excessive mucus accumulation and frequent inflammation (Ehre *et al*, 2014). We thus wanted to see whether the transcriptome alterations in mutant intestinal organoids translate into a cystic fibrosis-like phenotype. Staining mucin polysaccharides by periodic acid-Schiff (PAS) showed intense mucus accumulation in goblet cells specifically in *Cdk8*^*-/-*^ *Cdk19*^*-/-*^ organoids (Fig 4A, B). Time-lapse video-microscopy confirmed accelerated mucus release from double mutant organoids (Fig 4C, D). Functionality of CFTR can be tested in intestinal organoids using forskolin, an adenylate cyclase activator that, if CFTR is functional, induces luminal fluid secretion and organoid swelling (Dekkers *et al*, 2013). We found that forskolin caused swelling of wild-type but not double mutant organoids (Fig 4E, F; Movie S1), indicating that CFTR downregulation upon loss of CDK8 and CDK19 functionally recapitulates the *Cftr* mutant phenotype.

**Fig. 4.**
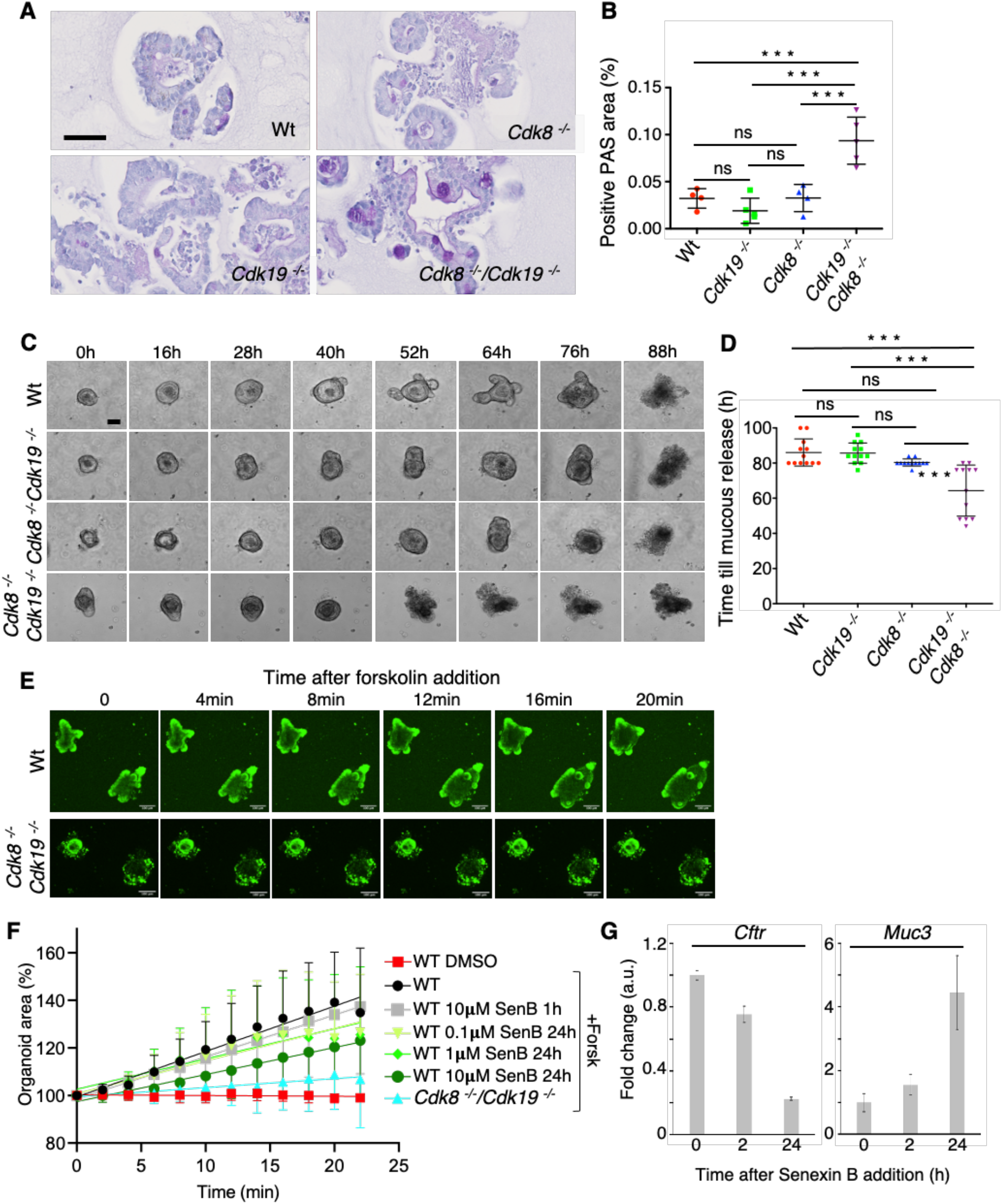
CDK8 and CDK19 regulate the CFTR pathway in the small intestine. **(A)** Histological PAS staining of organoids treated for 7 days with OH-tamoxifen. Scale bar, 50μm. **(B)** Quantification of PAS signal (% of total organoid area; mean ± SD are shown) in the four different genotypes presented in (A). Areas containing positive PAS staining were detected and quantified using QuPath and ImageJ programs. Adjusted p-values of ordinary one-way Anova followed by Tukey’s multiple comparison test are indicated: (***) p-value ≤ 0.001; ns: not significant (p > 0.05). **(C)** Representative phase contrast images of organoids at the indicated time points after 7 days of OH-tamoxifen treatment are shown. Scale bar, 100 μm. **(D)** Quantification of the time needed for mucus release (observed as a dark staining in the center of the organoid; mean ± SD are shown). Adjusted p-values of ordinary one-way Anova followed by Tukey’s multiple comparison test are indicated: (***) p-value ≤ 0.001; ns: not significant (p > 0.05); (n= 17 for Wt, n=11 for *Cdk8* ^*-/-*^, n=18 for *Cdk19* ^*-/-*^, n= 12 for *Cdk8* ^*-/-*^*/ Cdk19* ^*-/-*^). **(E)** Fluorescence confocal microscopy images of Calcein green–labeled WT and Cdk8^-/-^/Cdk19^-/-^ organoids treated with forskolin. Scale bars, 100 μm. **(F)** Quantification of forskolin-induced swelling in WT organoids treated for 1hour or 24 hours with 0.1μM, 1μM or 10μM Senexin B (SenB), as indicated, or double KO organoids; DMSO vehicle was used as control. The surface area of individual organoids at different time points relative to the area at t = 0 (100%) was measured (mean ± SD, n=8). Linear regression lines are shown. **(G)** qRT-PCR analysis of Cftr and Muc3 mRNA levels in WT organoids either not treated (t=0), or treated with 10µM Senexin B for 2 or 24 hrs.

These results implicate CDK8 and CDK19 as functionally redundant regulators of the CFTR pathway in the small intestine. Since transcriptional regulation by the Mediator CKM module might be in part independent of CDK8/19 kinase activity, we investigated whether inhibiting CDK8/19 would recapitulate their genetic disruption. We treated wild-type organoids with the CDK8/CDK19 inhibitor, Senexin B, for 1h or 24h before adding forskolin. We reasoned that if effects of CDK8/19 inhibition depend on transcriptional changes, they might take 24h to become detectable, whereas if they depend only on post-transcriptional regulation of CFTR, they might be seen after 1h. Fig 4F shows that there is a dose-dependent reduction of swelling after 24h, but not 1 hour, of Senexin B treatment, implicating that loss of CDK8/19 kinase activity recapitulates a *Cftr-*mutant phenotype. To see whether this correlates with downregulation of CFTR expression, we performed qRT-PCR analysis on organoids treated with Senexin B over a time course. We found that Senexin B treatment for 24h leads to the downregulation of *Cftr* and upregulation of *Muc3* expression (Fig 4G) seen upon genetic ablation of both *Cdk8* and *Cdk19*, indicating that kinase activity of CDK8/19 controls CFTR pathway gene expression.

This study shows that Mediator kinases are both functionally redundant and largely dispensable for cell survival, proliferation, and differentiation, but may be essential for regulation of specific gene sets in particular cell types; in this case, the CFTR pathway in the intestinal epithelium. Nevertheless, while cell proliferation defects have not previously been reported in HCT-116 cancer cells lacking both kinases (Koehler *et al*, 2016) our data suggest that in the intestinal epithelium, cells devoid of both CDK8 and CDK19 have an increased tendency to become quiescent, implying that they provide a growth advantage.

Our *in vivo* results do not support an oncogenic role for CDK8 in intestinal tumourigenesis, in contrast to early *in vitro* studies (Firestein *et al*, 2008; Morris *et al*, 2008). Another recent study using the heterozygous germline *Apc*^*min*^ mutant model of intestinal tumourigenesis also concluded that *Cdk8* deletion does not hinder tumourigenesis; on the contrary, in this model, while there was no difference in micro-adenoma formation, detectable increases in tumour number, size and fraction of proliferating cells were observed upon deletion of *Cdk8* (McCleland *et al*, 2015). The reasons for the slight difference in effects of *Cdk8* deletion between chemical carcinogenesis and *Apc*^*min*^ mutation on tumours are currently unclear, but, taken together, these studies suffice to conclude that *Cdk8* has neither oncogenic nor strong tumour suppressor activity in the mouse intestine.

Our results also do not support an essential role for Mediator kinases in general gene expression, in contrast to Mediator itself, since relatively few genes were highly deregulated in double knockouts. Slightly more genes were upregulated than downregulated upon loss of either CDK8 alone or both kinases, while effects of combined deletion were more than additive of effects of single deletions, indicating functional redundancy. Organoid growth and differentiation were not prevented by knockout of both kinases, indicating that, generally, CDK8 and CDK19 are not essential for implementation of new transcriptional programmes. However, we found that there was a strong overlap between transcriptome changes of double knockout organoids and intestinal knockout of the gene encoding CFTR, a chloride and bicarbonate ion-channel that regulates fluid homeostasis in epithelia, and whose mutation causes cystic fibrosis (CF), a disease associated with mucus retention and inflammation of epithelia. Double knockout organoids showed increased mucin expression and strong accumulation of mucins in goblet cells, coupled with a precocious secretion of mucus, as well as a lack of forskolin-induced swelling, which depends on CFTR (Dekkers *et al*, 2013), indicating that CDK8/19 regulate fluid and/or mucus homeostasis. This appears to depend on their kinase activity, as specific inhibition of both kinases for 24 hours using Senexin B impaired forskolin-induced swelling in a dose-dependent manner. Since acute CDK8/19 inhibition in organoids for one hour prior to the forskolin assay had no effect, this appears to be due to transcriptional downregulation of *Cftr*. Whether or not CDK8 or CDK19 are implicated in the pathogenesis of cystic fibrosis, however, remains an open question. Almost all CF patients harbour genetic mutations in the *CFTR* gene, yet identical mutations do not have identical disease severity, and the variability between patients is associated with different genetic loci (Wright *et al*, 2011; Corvol *et al*, 2015). There are also significant but variable gene expression alterations in *CFTR* mutant cells and upon therapeutic interventions, some of which may influence disease phenotypes (Hodos *et al*, 2020). The genes encoding CDK8, CDK19 and cyclin C have not so far been associated with CF. *CDK19* is downregulated upon several model therapeutic interventions, including overexpression of the micro-RNA miR-138, which promotes CFTR expression (Hodos *et al*, 2020; Ramachandran *et al*, 2012). However, in our study, CDK19 knockout alone was insufficient to cause a CF phenotype in the intestine, suggesting that variation in CDK19 expression does not affect CFTR. Identifying the mechanisms by which CDK8 and CDK19 affect expression of genes in the CFTR pathway will be important to better understand the pathophysiology of cystic fibrosis, but will require further studies.

## Materials and Methods

### Animal studies

All animal experiments were performed in accordance with international ethics standards and were subjected to approval by the Animal Experimentation Ethics Committee of Languedoc Roussillon.

### Cdk8 *conditional knockout mice*

*Cdk8*^*lox/lox*^ mice were generated as follows: An 8076 bp genomic fragment (mouse chromosome 5: 146,254,503 to 146,262,579) enclosing the essential exon 2 (whose deletion results in loss of the essential catalytic lysine residue and causes a frameshift truncating over 90% of the protein) of the CDK8 gene was amplified by PCR from genomic DNA of 129/Sv embryonic stem cells and cloned into pGEM-T-easy. The diphtheria toxin A gene was cloned into the SacII site. 64 bp to the 3’ of exon 2, the sequence CTCTAT was mutated to CTCGAG, generating an XhoI site. LoxP sites flanking exon 2 were generated by a combination of conventional cloning and recombineering, using a recombineering approach (Liu *et al*, 2003). The loxP PGK-Neo cassette was amplified from pL452 plasmid with flanking AvrII/HindIII sites at each end and cloned into the AvrII site upstream of exon 2. Fragment orientation was confirmed by the generation of 3.5 kb HindIII and 2.0 kb NheI sites, and the vector was recombined in *E. coli* strain SW106 with inducible Cre recombinase expression followed by HindII digestion, generating a single loxP site upstream of exon 2. Into this recombined vector, the FRT-PGK-Neo-FRT-LoxP cassette (amplified from pL451 with flanking XhoI sites) was cloned in the newly generated XhoI site downstream of exon 2, resulting in the “deletion construct”. The orientation was confirmed by the generation of 2.2 kb Nhe1 and 3.4 kb BamHI sites. Functionality of the two recombination sites was tested as follows: the FRT site was confirmed by recombination in *E. coli* strain SW105 with inducible FlpE recombinase expression, deleting the FRT-Neo cassette and generating a 1.4 kb BamHI fragment; the resulting plasmid was transformed in *E. coli* strain SW106 with inducible Cre recombinase expression, deleting exon 2 and resulting in a 1.1 kb BamHI fragment. The NotI linearised fragment of the deletion construct was transfected by electroporation into 129/Sv embryonic stem cells. 244 Neomycin-resistant colonies were genotyped by PCR and Southern blotting. Two probes were used: one outside the 3’ end of the deletion construct, with HindIII digestion site giving a single 9kb fragment for the WT and a 7kb fragment for the correctly-integrated deletion cassette, and one to the 5’ end of the deletion cassette, again giving the same 9kb fragment for the Wt but a 3.5 kb fragment for the deletion cassette. 10 colonies showed a correct integration by homologous recombination. These ES cells were injected into blastocysts obtained from pregnant BALB/C mice, and chimeric mice were crossed with C57/Bl6J mice constitutively expressing FlpE recombinase, removing the FRT-Neo cassette. Agouti mice were genotyped by PCR, showing correct insertion of the LoxP sites around exon 2.

*Cdk8* ^*lox/lox*^ mice were crossed with *Villin-Cre-*^ERT2 +/-^ mice to obtain *Cdk8* ^*lox/lox*^, *Villin-Cre-*^ERT2 +/-^.

### Tamoxifen treatment of mice to induce Lox recombination

Mice were first injected intraperitoneally (IP) with 100μl of 20mg/ml tamoxifen solution (in corn oil). After the injection, they were fed during 5 days with cookies containing 400mg tamoxifen citrate per kg diet (Envigo, Ref TD.130859).

### Apc/Cdk8 conditional knockout mice

C57BL/6 *Apc* ^*lox/lox*^ mice (Colnot *et al*, 2004) were provided by Philippe Jay (IGF, Montpellier). These mice were crossed with *Cdk8* ^*lox/lox*^/*Villin-Cre-*^*ERT2 +/-*^ to obtain *Apc* ^*lox/lox*^*/Cdk8* ^*lox/lox*^/*Villin-Cre*-^ERT2 +/-^ mice. IP injection with tamoxifen during 5 days induced *Cdk8* exon 2 deletion and *Apc* exon 14 deletion in the intestines of mice containing the *Villin-Cre-*^*ERT2*^ gene. Small intestine and colon samples from these mice were genotyped and analysed by IHC and Western blotting.

### Cdk7/Cdk8 conditional knockout mice

Cdk8^lox/lox^ mice were crossed with RERT mutant mice expressing the inducible Cre-ERT2 from the endogenous *Polr2a* locus (Guerra *et al*, 2003). The Cdk8^lox/lox^ RERT mice were then crossed with Cdk7^lox/lox^ mice (Ganuza *et al*, 2012) to obtain Cdk8^lox/lox^/Cdk7^lox/lox^, RERT mice in which CDK8 and CDK7 proteins should be removed from the whole body after tamoxifen treatment. Animals were sacrificed after tamoxifen treatment and intestinal epithelium was collected as indicated in the Sample preparation section below. Proteins were extracted and analysed by Western blotting.

### AOM/DSS-induced colon carcinogenesis

11 *Cdk8* ^*lox/lox*^ and 11 *Cdk8* ^*lox/lox*^/*Villin-Cre-*^ERT2 +/-^ mice were treated with tamoxifen as described above. 4 days later, mice (*Cdk8* ^*lox/lox*^ and *Cdk8* ^*-/-*^) were given a single intraperitoneal injection of AOM (10mg/kg in 0.9% saline; A5486, Sigma-Aldrich); 5 days later, 2.5% Dextran Sodium Sulfate (DSS; MP Biomedicals) was administered in the drinking water during 5 consecutive days. DSS treatment was repeated two more times with 16 days intervals without DSS for recovery (see scheme, Fig S4A). Mice were sacrificed 16 days after the third DSS treatment. Colons were flushed with PBS and either used for intestinal epithelium extraction (see Sample preparation for details) or used for IHC studies. Colons used for IHC were fixed overnight in neutral buffered formalin (10%) before paraffin embedding. Briefly, 4μm thick sections were dewaxed in xylene and rehydrated in graded alcohol baths. Slides were incubated in 3% H_2_O_2_ for 20 min and washed in PBS to quench endogenous peroxidase activity. Antigen retrieval was performed by boiling slides for 20 min in 10mM sodium citrate buffer, pH 6.0. Nonspecific binding sites were blocked in blocking buffer (TBS, pH 7.4, 5% dried milk, 0.5% Triton X-100) for 60 min at RT. Sections were incubated with anti-β-catenin antibody diluted in blocking buffer overnight at 4°C. Envision+(Dako) was used as a secondary reagent. Signals were developed with Fast DAB (Sigma-Aldrich). After dehydration, sections were mounted in Pertex (Histolab), imaged using the Nanozoomer-XR Digital slide Scanner C12000-01 (Hamamatsu) and analysed using NDP.view 2 program (Hamamatsu).

### Small intestine organoids

*Cdk8* ^*lox/lox*^ and *Cdk8* ^*lox/lox*^/*Villin-Cre-*^*ERT2*^ mice were used to obtain small intestine organoids. Establishment, expansion and maintenance of organoids were performed as described previously (Sato *et al*, 2009). To induce the Cre-mediated recombination of *Cdk8 in vitro*, organoids were cultured during 7 days in medium supplemented with 600nM 4-Hydroxytamoxifen (Sigma H7904) resuspended in ethanol. Evaluation of knockout efficiency was performed using genotyping, qPCR and Western blotting.

CRISPR/Cas9-mediated genome editing was employed to remove CDK19 from the organoids. CRISPR single guide RNA (sgRNA) targeting murine *Cdk19* sequence (5’-AAAGTGGGACGCGGCACCTA-3’, from Zhang lab database) was cloned as synthetic dsDNA into lentiCRISPRv2 vector as described ((Sanjana *et al*, 2014); provided by F. Zhang, Addgene plasmid #52961). Lentiviruses encoding the sgRNA targeting sequence were produced in HEK 293T cells transfected with LentiCRISPRv2 (+sgRNA *Cdk19*), pMD2.G and psPAX2. The viral supernatant (collected in organoids culture media) was passed through a 0.45-μm filter and used the same day for infection. Lentiviral-mediated transduction and antibiotic selection was performed as described previously (Onuma *et al*, 2013). Briefly, for lentiviral infection, organoids (5 days after seeding) were diluted into 10ml of PBS and dissociated into single cells by passing them 10-15 times through a needle with an insulin syringe. A volume containing 1–5 × 10^5^ intestinal cells was centrifuged at 300g for 5 minutes and resuspended with 1ml of the viral supernatant produced in HEK 293T cells. This mixture (virus + single stem cells) was layered on top of a Matrigel-covered well (12 well plate). 24 hours later, virus and dead cells containing media were removed and the Matrigel-attached cells were covered with 200μl of Matrigel + 200μl of culture medium to create a “sandwich” containing the infected cells inside. After polymerisation of the second Matrigel layer, 1 ml of organoid media per well was used to allow organoid formation inside the Matrigel. 24 hours later, Puromycin was added (5μg/mL) and selection was conducted for 4 days. Once the organoids appeared (4-5 days after seeding the infected single cells), CDK19 knockout was verified by Western blot. We observed that CDK19 protein was still present, albeit decreased; therefore, we picked individual organoids and allowed them to growth in separated wells until we obtained several populations where CDK19 protein was completely absent, as seen by Western blot. DNA sequencing confirmed the deletion of a fragment of DNA around the sequence corresponding to *Cdk19* sgRNA, and qPCR confirmed the absence of *Cdk19* mRNA.

### Mouse intestine epithelium, colon tumour and organoid sample preparation

For intestine epithelium samples, a fragment of intestine was cut and flushed with PBS. It was incubated for 5-10 minutes in EDTA-containing buffer (500ml RPMI, 20mM Hepes, 1% Penicillin/Streptomycin (Sigma P4333), 12.5μg/ml DTT, 2mM EDTA pH 7.4 and 10% FBS) to allow easy detachment of the intestine epithelium. The intestinal tube was opened longitudinally and put over a horizontal plate to allow scrapping of the epithelial cells by trawling two needles in opposite directions over the tissue. Cells were recovered from the plate by wetting them with a small volume (200μl) of PBS. They were put into an Eppendorf tube where they were spun down to remove most of the PBS. Pellets were snap frozen and conserved at -80°C. Colon tumours were cut from the colon, washed with PBS and frozen in liquid nitrogen.

Frozen intestinal epithelium and colon tumour samples were resuspended in 250μl of lysis buffer with protease and phosphatase inhibitors (5mM Tris, pH 7.4, 100mM NaCl, 50mM NaF, 40mM beta-glycerophosphate, 2.5mM Na-Vanadate, 5mM EDTA, 1mM EGTA, 1mM DTT, 1% Triton X-100 and Protease inhibitor cocktail diluted 1/400 (Sigma P8340)) and, after addition of 100μl of stainless-steel beads (0.2 mm diameter, 1lb, Next Advance, SSB02), they were disrupted in a bullet blender storm 24 (Next Advance) by shaking during 4 minutes at 4°C with an intensity level of 8. The lysate was incubated during 20 more minutes at 4°C (without shaking) and the solubilised proteins were recovered from the supernatant by centrifugation at 16000g for 20 minutes at 4°C, frozen in liquid nitrogen and stored at -80°C until use. Protein concentrations were determined by BCA protein assay (Pierce Biotechnology).

For organoids samples, Matrigel was disrupted by pipetting up and down several times the media in each well over the dome of Matrigel. This mix was spun down at 200g for 5 minutes at 4°C and the pellet was washed twice with 1ml of PBS. Pellets were snap frozen in liquid nitrogen and kept at -80°C until use.

For organoids extracts, frozen pellets were lysed by incubation at 4°C for 20 minutes in lysis buffer with protease and phosphatase inhibitors (50mM Tris, pH 7.4, 100mM NaCl, 50mM NaF, 40mM beta-glycerophosphate, 2.5mM Na-Vanadate, 5mM EDTA, 1mM EGTA, 1mM DTT, 1% Triton X-100 and Protease inhibitor cocktail (Sigma P8340) diluted 1/400. The solubilised proteins were recovered from the supernatant after centrifugation at 16000g for 20 minutes at 4°C, snap frozen in liquid nitrogen and stored at -80°C until use. Protein concentrations were determined by BCA protein assay (Pierce Biotechnology).

### Western-blotting

For intestinal epithelium, colon tumours and organoids, 30μg of of total proteins were separated by SDS-polyacrylamide gel electrophoresis (SDS-PAGE) (7% 10% and 12.5% gels) and transferred to Immobilon membranes (Milipore) at 90 volts for 120 min with a wet-blotting apparatus. Membranes were blocked in TBS-T pH 7.6 (20mM Tris, 140mM NaCl, 0.1% Tween-20) containing non-fat dry milk (5%), incubated with the primary antibody in TBS-T + 3% BSA for 2 hours at RT or over-night at 4°C, washed 3 times with TBS-T for a total of 15 minutes, incubated with secondary antibody at 1/10000 dilution in TBS-T + 5% nonfat dried milk for 30 minutes at RT, and washed 3 times in TBS-T for a total of 15 minutes. Signals were detected using Western Lightning Plus-ECL (PerkinElmer) and Amersham Hyperfilm^™^ECL (GE Healthcare).

### RNA isolation and qRT-PCR

RNA was extracted from organoids and purified using RNeasy mini kit (Qiagen) according to manufacturer’s protocol. For reverse transcription (cDNA synthesis), 1μg of purified RNA in total volume of 13µl, extracted by RNeasy Mini Kit (Qiagen), was mixed with 1µl of 10mM dNTPs mix (LifeTechnologies) and 1µl of 10µM oligo(dT) 20-primer. Samples were incubated at 65°C for 5 minutes, transferred to ice. 4µl of 5x First Strand Buffer, 1µL of 100mM DTT, 1µl of RNase OUT RNase Inhibitor (Invitrogen) and 1µl of SuperScript® III Reverse Transcriptase (LifeTechnologies) were added to each sample and incubated at 50°C for 1 hour. The reaction was inactivated at 70°C for 15 minutes.

qPCR was performed using LightCycler® 480 II (Roche). The reaction contained 5μl of a 1/10 dilution of the cDNA obtained after RT (25ng of cDNA), 1µl of 10μM qPCR primer pair, 10µl 2x Master Mix in a final volume made up to 20µl with DNase free water. qPCR was conducted at 95°C for 5 min, followed by 45 cycles of 95°C 10s, 59°C 20s and 72°C 15s. The specificity of the reaction was verified by melting curve analysis. β2-mioglobulin (B2M) was used as housekeeping gene.

### Genomic DNA extraction and genotyping

For colon tumors, genomic DNA was extracted using the KAPA Mouse genotyping kit from Clinisciences (KK7352) directly over 3-4 tumors that had been removed from the colon before the extraction of intestinal epithelium.

Genomic DNA was extracted from intestinal epithelium pellets or organoids using KAPA Mouse genotyping kit from Clinisciences (KK7352).

The forward and revers primers used for amplification of the *Cdk8* wild-type allele (650 bp), the *Cdk8*^*Lox/Lox*^ (850 bp) and the fragment obtained after tamoxifen-induced Cre recombination of *Cdk8*^*Lox/Lox*^ (*Cdk8* Lox + Cre (-)) (340 bp) are indicated in the table below as CDK8 Fw (genotyping) and CDK8 Rev (genotyping). These fragments were amplified from genomic DNA using the following PCR protocol: 95°C for 5 min; 35 cycles at 95°C for 15 s; 57°C for 15 s; 72°C for 1min 30 s. Amplified DNA fragments were migrated in a 1,5% agarose gels and stained with Ethidium bromide for detection.

### Primers used for qPCR and genotyping

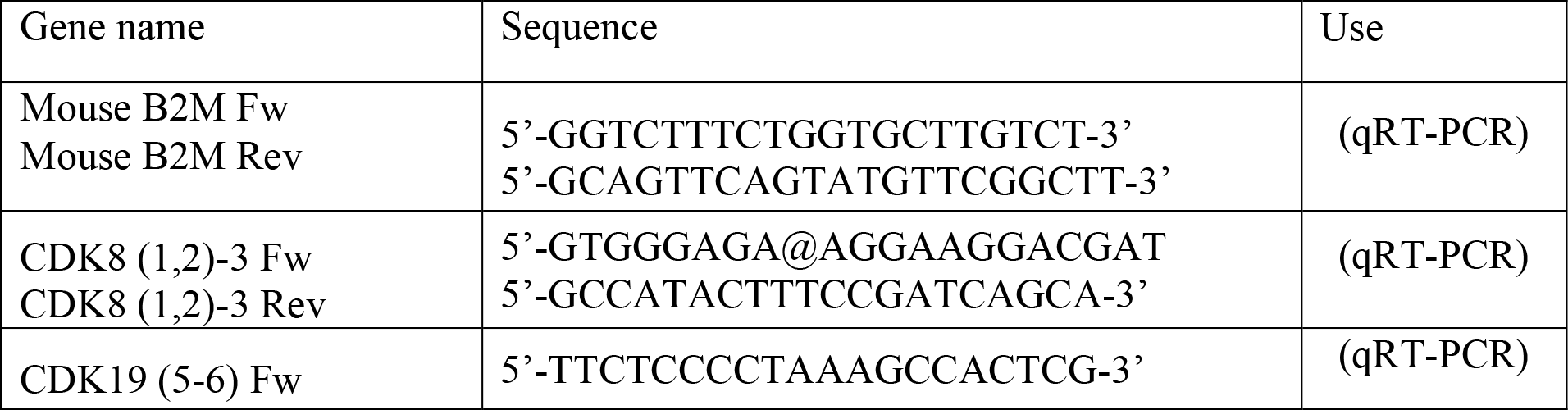
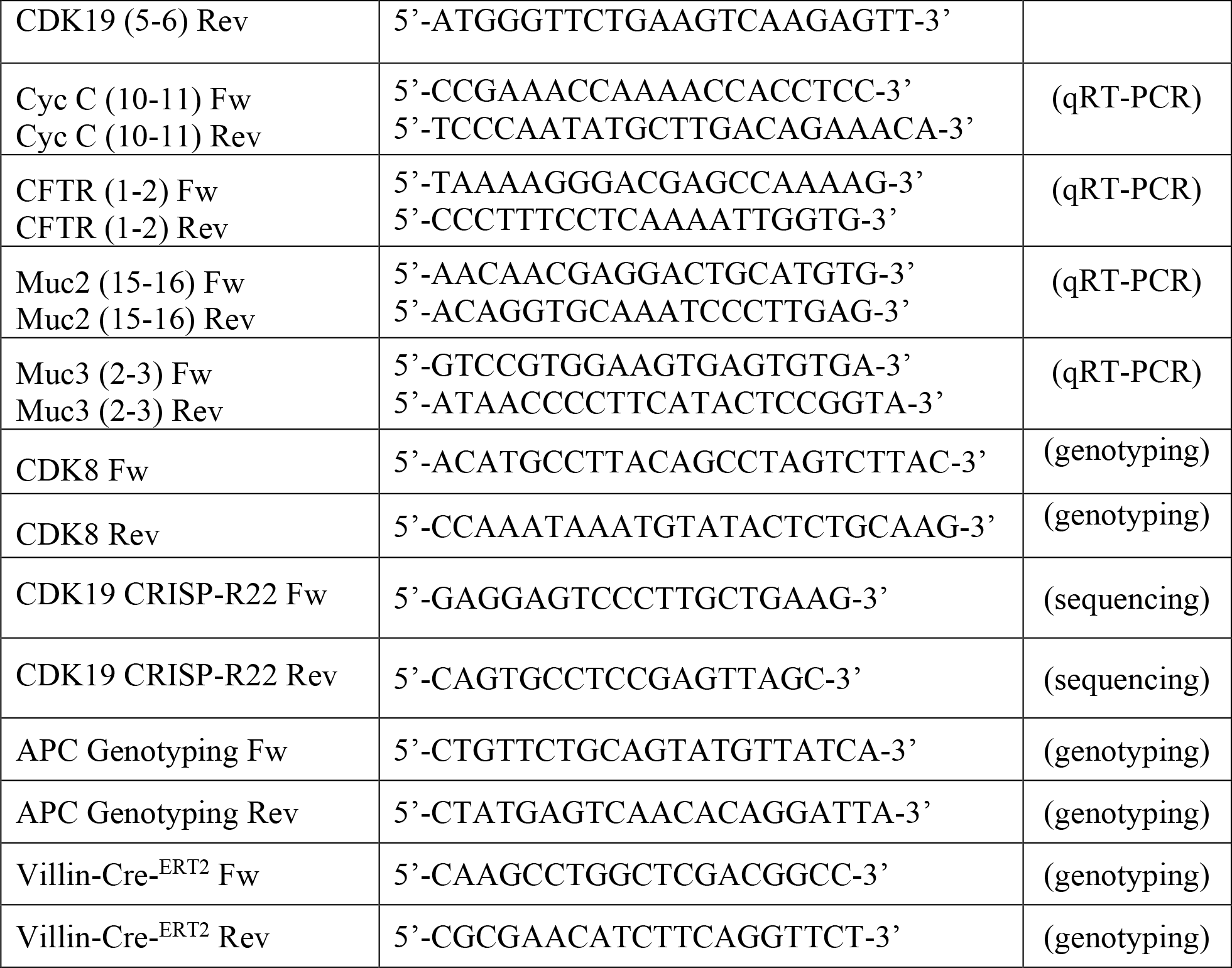

### Antibodies

*Antibodies.*
| Name | Clone | Source | Species | Cat# |
| --- | --- | --- | --- | --- |
| CDK8 (C19) | Polyclonal | Santa Cruz | Goat | sc-1521 |
| CDK19 | Polyclonal | Sigma | Rabbit | HPA007053 |
| Cyclin C | Polyclonal | gift | Rabbit |  |
| PCNA | Monoclonal | Santa Cruz | Mouse | sc-56 |
| Lysozyme | Polyclonal | Dako | Rabbit | A0099 |
| Anti-Dclk1 | Polyclonal | Abcam | Rabbit | Ab31704 |
| Olfm4 | Monoclonal | Cell signaling | Rabbit | 39141 |
| β-Catenin | Monoclonal | BD<br>BioSciences | Mouse | 610154 |
| BrdU | Monoclonal | DSHB | Mouse | G3G4 |
| RNA pol II<br>Ser2P | Monoclonal | JC Andrau's<br>lab | Rat | (3E10) |
| RNA pol II<br>Ser5P | Monoclonal | JC Andrau's<br>lab | Rat | (3E8) |
| RNA pol II<br>Ser7P | Monoclonal | JC Andrau's<br>lab | Rat | (4E12) |
| CDK7 | Monoclonal | Santa Cruz | Mouse | sc-7344 |
| Ki-67 | SP6 | Spring<br>Bioscience | Rabbit | M3064 |
| Ki-67 | Monoclonal | Abcam | Rabbit | Ab16667 |
| p53 | CM5 | Novocastra | Rabbit | NCL-p53-CM5p |
| Cleaved Caspase 3 | ASP176 | Cell signaling | Rabbit | 9661S |
| Phospho-Stat1 (Ser727) | Polyclonal | Cell signaling | Rabbit | #9177 |
| CFTR | Monoclonal | Abcam | Mouse | ab2784 |
| β-Actin | Monoclonal | Sigma | Mouse | a5441 |
| GAPDH | Polyclonal | Sigma | Rabbit | G9545 |

CycC purified antibody. Rabbit anti-Cyclin C serum was a kind gift from Jacques Piette (IGMM Montpellier, France (Barette *et al*, 2001)). Cyclin C specific antibodies were purified from serum by incubation with a membrane containing Cyclin C protein (ProQinase GmbH). The antibodies were eluted with 200μl of 0,2M glycine pH 2.5 and neutralised rapidly with 21μl of 1M Tris. RNA Pol II Ser2P (3E10), RNA Pol II Ser5P (3E8) and RNA Pol II Ser7P (4E12) antibodies were a kind gift from Jean-Christophe Andrau’s lab (IGMM, CNRS Montpellier, France;(Chapman *et al*, 2007)).

### Immunohistochemistry (IHC) and tissue staining

Whole intestines were flushed with PBS and turned inside-out on a wooden stick. They were collected and fixed 24h in neutral buffered formalin 10%, dehydrated, and embedded in paraffin.

Organoids were collected and fixed for 1h in 4% paraformaldehyde at room temperature. They were washed with PBS (x2) and resuspended into 100µl of Histogel (Fisher Scientific, Ref 12006679), previously thawed in a hot water bath at 60°C. Each drop containing the organoids and Histogel was dried on top of a flat surface and embebed in paraffin. Paraffin-embedded intestines or organoids were cut into 3-μm-thick sections, mounted on slides, then dried at 37°C overnight. Tissue sections were stained with hematoxylin-eosin (HE) with HMS 740 autostainer (MM France) for preliminary analysis. For mucosubstances, tissue sections were stained with Periodic Acid Schiff’s (PAS) staining (J. Bancroft & A. Stevens, 1982).

For Ki67, p53 and Caspase 3, IHC was performed as described previously (Rahmanzadeh *et al*, 2007), on a VENTANA Discovery Ultra automated staining instrument (Ventana Medical Systems), using VENTANA reagents, according to the manufacturer’s instructions. Briefly, slides were de-paraffinised, then epitope retrieval was performed with CC1 solution (cat# 950-124) at high temperature (95-100°C) or for CC2 solution (cat# 950-123) at high temperature (91°C) for a period time that is suitable for each specific antibody. Endogenous peroxidases were blocked with Discovery Inhibitor (cat# 760-4840).

For CDK8 immunostaining, signal enhancement was performed using the rabbit antibody anti-goat (Vector Laboratories, cat#BA-5000, 1:2000,) for 16min at 37°C then using DISCOVERY OmniMap anti-rabbit HRP detection Kit (cat# 05269679001) according to the manufacturer’s instructions. The same kit was used for all other rabbit primary antibodies to amplify the signal. Slides were incubated with DAB (cat# 05266645001), then counterstained with hematoxylin II (cat# 790-2208) for 8 min, followed by Bluing reagent (cat# 760-2037) for 4 min. Slides were then dehydrated with Leica autostainer and coverslipped with Pertex mounting medium with CM6 coverslipper (Microm). Brightfield stained slides were digitalised with a Hamamatsu NanoZoomer 2.0-HT scanner and images were visualised with the NDP.view 1.2.47 software.

For Paneth cells, tuft cells, β-catenin and Brdu detection, the IHCs were performed manually. After deparaffination and rehydration, demasking of antigenic sites was performed by boiling the slides for 20’ in 10mM Na-Citrate pH 6.4. After cooling down, slides were treated with 3% H_2_O_2_, 5’ at RT for peroxidase inhibition. Samples were blocked in blocking solution (TBS, 0.5% Triton, 5% dry milk) for 20’. First antibody was diluted 1/400 in blocking solution and the slides were incubated O/N at 4°C in a humid chamber. Slides were then washed with TBS + 0.1% Tween 20 (3 times) and once with TBS without Tween. Secondary antibodies (ImmPRESS^™^reagent kit peroxidase anti-rabbit (MP-7451) or anti-Mouse (MP-7402), from Vector Laboratories) were incubated for 30’ at RT. Slides were washed twice with TBS + 0.1% Tween 20 and the final wash was done in H_2_O. Peroxidase staining was performed using Sigma Fast DAB tablet set (D4293-50SET). After Hematoxylin staining of the nucleus with Gill’s Hematoxylin solution N°2 (CAS 517-28-2) from Santa Cruz (SC-24973), slides were rehydrated and mounted in Pertex^R^ (Histolab 00811).

### Sequence alignment

CDK8 and CDK19 protein sequences from different species were aligned using the BoxShade server (https://embnet.vital-it.ch/software/BOX_form.html).

### RNA sequencing

After 7 days of treatment with 600nM 4-Hydroxytamoxifen, organoids were lysed and RNA was extracted following the Trizol RNA isolation protocol (W.M. Keck Foundation Biotechnology Microarryay Resource Laboratory at Yale University) until the end of the phase-separation step. Total RNA cleanup with DNase digestion was performed by addition of 1.5 volumes of absolute ethanol on top of the aqueous phase obtained after the phase-separation and following the Qiagen RNeasy protocol (W.M. Keck Foundation Biotechnology Microarray Resource Laboratory at Yale University). RNA integrity was analysed on Agilent 2100 bioanalyzer. All conditions were prepared as biological triplicates and sent to BGI Tech Solutions (Hong Kong)-Co for library preparation and RNA sequencing. Purification of mRNA from total RNA was achieved using oligo(dT)-attached magnetic beads and then fragmented for random hexamer-primed reverse transcription, followed by a second-strand cDNA synthesis. Sequencing was performed with the BGISEQ, SE50 platform to obtain an average of 50 million single-end, 50bp reads per sample.

### Bioinformatic analysis

The raw reads obtained in fastq format were subject to quality control using the FastQC software. The reads passing the quality control were aligned to the mouse reference genome (GRCm38.p6) and the counts per gene were quantified using the tool STAR 2.6.0a(2). The Ensembl mouse genome annotations (release 93) were used to map the reads to each gene and their corresponding transcripts. Differential gene expression analysis was performed in R using the DESeq2 library. MA-plots showing the deferentially expressed genes were generated with an in-house script and the gene set enrichment analysis were performed using the “enrichR” library, in both cases using the R programming language.

### *CRISPR Cas9-targeting of* Cdk19 *gene*

CRISPR single guide RNA (sgRNA) targeting exon 1 in mouse *Cdk19* (CDK19_Mouse: 5’-AAAGTGGGACGCGGCACCTA-3’ from Zhang lab database) was cloned as synthetic dsDNA into lentiCRISPRv2 vector (provided by F. Zhang, Addgene plasmid #52961) as described (Sanjana et al., 2014). Generation of lentiviral particles and infection of organoids were carried out following classical procedures, as described previously (Onuma *et al*., 2013). Successfully infected cells were selected with puromycin (5μg/ml) during 4 days. Single organoids were picked for clonal expansion. Effects of targeted deletion were verified by sequencing. Genomic DNA was extracted from organoids using KAPA Mouse genotyping kit from Clinisciences (KK7352). The 2 primers used for amplification of the *Cdk19* allele are indicated in the table above as CDK19 CRISP-R22 Fw and CDK19 CRISP-R22 Rev (sequencing). They were amplified as follows: 95°C for 5 min; 30 cycles at 95°C for 45’’; 57°C for 30’’; 72°C for 1min 30 s. Absence of CDK19 protein in the organoids was also verified by Western blotting.

### Time-lapse microscopy

Organoids (Wt, *Cdk8*^*Lox/Lox*^, *Cdk19*^*-/-*^ *and Cdk8*^*Lox/Lox*^*/Cdk19*^*-/-*^) were treated with 600nM 4-hydroxytamoxifen during 7 days to induce recombination of the LoxP sites flanking *Cdk8* exon 2. Images were taken every 4 hours during 7 days on an inverted microscope (Axio Observer, Zeiss) equipped with a heated chamber allowing constant temperature (37°C) and CO_2_ flow (5% CO_2_). CCD camera (Princeton Instruments (Micromax), pixel = 6,7 μm), with 10x/0.3 DRY PH1objective, correction ECPLAN Neofluar, 5.2 mm working distance.

Acquisition software was MetaMorph 7.8 (Molecular Devices, LLC). Images were analysed using Image J software to calculate the time for release of mucus in the lumen of the *organoid* (observed as a dark staining in the center of the organoid).

### Forskolin-induced swelling

To remove exon 2 of *Cdk8* from *Cdk8*^*lox/lox*^/Villin-Cre-ERT2^+/-^/*Cdk19*^-/-^, organoids were treated with 600nM 4-hydrxoytamoxifen for 7 days. Once the *Cdk8/Cdk19* double KO was obtained, forskolin-induced swelling was measured as indicated (Dekkers *et al*, 2013). Organoids were transferred to CELLview culture dishes PS 35/10 mm, glass bottom, 4 compartments (Greiner Bio-One, 627870), two days before imaging. Confocal spinning disk (Dragonfly, Andor, Oxford Instruments) microscope equipped with heated chamber allowing constant temperature (37°C) and CO_2_ flow (5% CO_2_), EMCCD iXon888 camera (Lifer Andor, pixel = 13 μm), objective 10x/0.45 DRY, correction Plan Apo Lambda, 4mm working distance, was used for imaging, with Fusion acquisition software. Images of a single organoid, previously selected, were taken every 2 minutes during 20 minutes after forskolin addition (5µM, or DMSO vehicle control) to the media. For data analysis, a macro was created using Fiji software. It consisted of recognising and filling the structures imaged through the alexa-488 track, to calculate the increase of total organoid area in single organoids over the different time points.

### Statistics

Graphs and statistical analyses were performed using Microsoft Excel 16.50 and GraphPad Prism6 using analyses described in legends.

### Data availability

The RNA-sequencing data have been deposited in the Gene Expression Omnibus (GEO, NCBI) repository, and are accessible through GEO Series accession number GSE186377.

## Acknowledgements

L.A. was funded by the Montpellier FHU Cancer programme and by the Fondation pour la Recherche Medicale; E.H. was funded by the Ligue Nationale Contre le Cancer (LNCC); G.D. was funded by the National Cancer Institute (INCa); S.P., C.B-P. and F.G. are CNRS employees; D.F., P.J. and L.K. are Inserm employees. This work was undertaken with support from INCa (PLBIO10-068 and PLBIO15-005), and the LNCC (EL2013.LNCC/DF and EL2018.LNCC/DF). RHEM histology facility was supported by SIRIC Montpellier Cancer Grant INCa_Inserm_DGOS_12553, the European regional development foundation and the Occitanie region (FEDER-FSE 2014-2020 Languedoc Roussillon). We thank Sylvain de Rossi and Volker Baecker of the Montpellier Resources in Imaging (MRI) platform for help with microscopy and analysis.

## Author contributions

D.F. conceived the study with assistance from P.J., S.P., E.S., L.K. and D.F. designed experiments. S.P., E.S., E.H., L.A., A.B.A, A.C., C.B-P., N.P. and F.G. performed experiments. G.D. performed bioinformatics analysis. S.P., G.D., P.J., L.K. and D.F. interpreted data. S.P., L.K. and D.F. wrote the paper.

## Figure legends

**Fig S1. Mouse *Cdk8* conditional knockout by Lox/Cre targeting of exon 2. (A)** Schematic representation of the strategy used for the generation of *Cdk8*^*Lox*^ alleles from a genomic fragment of mouse enclosing exon 2 of the *Cdk8* gene. See Materials and Methods section for details. (**B)** Southern blot analysis of genomic DNA obtained from mice carrying the *Cd8*^*LoxFrt*^ and *Cdk8*^*Lox*^ alleles. DNA was digested with HindIII and probed with 2 different probes (5’ and 3’), whose positions are shown in the scheme in (A). The length of the fragments obtained after HindIII digestion of the wild type and the recombinant alleles are indicated (see also the scheme in (A)). **(C)** Left, scheme representing the control plasmids (a, b, and c) for WT, floxed and recombined *Cdk8* exon 2. The position of the oligos (Fw and Rev) used for PCR amplification of genomic DNA and control plasmids is indicated. Right, genotyping of *Cdk8* exon 2 in the mouse intestinal epithelium. All mice were treated with OH-tamoxifen to induce recombination of the LoxP sites. The recombined fragment appears as a 340 bp band in the *Cdk8* ^*-/-*^ and *Cdk8* ^*+/-*^ mice (*VillinCre*^*ERT2*^ recombinase-positive), and is absent in the *Cdk8* ^*Lox/+*^ mouse that does not contain the *VillinCre*^*ERT2*^ gene.

**Fig S2. CDK8 is not required for cell proliferation nor differentiation in mouse intestine**. **(A)** Representative immunohistochemistry images of Fig 1B. Small intestine, proximal and distal colon samples were stained for BrdU, Ki-67 or PCNA antibodies. Goblet cells were detected with PAS staining. Scale bars: 100μm for small intestine and distal colon, 50μm for proximal colon. **(B)** WB analysis of mouse intestine epithelium showing the absence of CDK8 protein 2 months after OH-tamoxifen feeding. Mice 4, 5 and 6 did not have the *VillinCre*^*ERT2*^ gene; mice 1, 2 and 3 had the *VillinCre*^*ERT2*^. GAPDH protein was used as loading control.

**Fig S3. Effects of CDK8 and CDK7 knockout on RNA pol II CTD phosphorylation**. WB analysis of the indicated proteins in mouse intestinal epithelium **(A)** or liver **(B)** samples from WT and *Cdk7* ^*lox/lo*x^, *Cdk*8 ^*lox/lo*x^, *Rpb-Cre-Ert2* ^*KI/KI*^ mice after tamoxifen treatment. Amido-black staining was used as loading control.

**Fig S4. CDK8 loss does not affect chemically-induced intestinal carcinogenesis. (A)** Scheme showing the steps of the AOM/DSS carcinogenesis experiment. See Materials and Methods for more detailed information. **(B)** Graphs showing female (left; n=6 in both groups) and male (right; n=5 in both groups) weight evolution over 21 days following the last DSS treatment. **(C)** Genotyping after AOM/DSS treatment confirms the recombination and loss of *Cdk8* exon 2 in colon tumors from *Cdk8* ^*-/-*^ mice. Control plasmids (a and b), are described in Fig S1C. PCR amplification with *Villin-Cre*^*ERT2*^-specific primers confirms the presence of the *Cre*^*ERT2*^ recombinase gene. **(D)** WB with the same colon tumour samples presented in (C) confirm the disappearance of CDK8 protein in the *Cdk8* ^*-/-*^ mice. β-actin was used as the loading control.

**Fig S5. CDK8 deletion does not prevent Apc-loss-dependent tumourigenesis in mouse intestine. (A)** Genotyping confirms the loss of *Cdk8* exon 2 in intestine epithelium from *Apc* ^*- /-*^/*Cdk8* ^*-/-*^ mice. Control plasmids (a, b and c) are described in Fig S1-C. **(B)** WB with intestine epithelium samples from mice presented in (A) confirm the absence of CDK8 protein in *Cdk8* ^*-/-*^ mice. Amido-black staining was used as loading control. **(C)** IHC staining of CDK8 and Ki-67 in small intestine and colon samples from *Apc* ^*-/-*^/*Cdk8* ^*+/+*^ and *Apc* ^*-/-*^/*Cdk8* ^*-/-*^ mice. Scale bars, 100μm. **(D)** Quantification of the Ki-67 positive area (% of the total area of the intestine presenting positive staining, quantified using QuPath (Bankhead *et al*, 2017) and Image J software; mean ± SD) in the IHC shown in (C). Ki-67 (n=16). Two-tailed p-value of unpaired t-test is indicated: ns, not significant (p > 0.05).

**Fig S6. Amino acid sequence conservation of CDK8 and its paralogue CDK19 between different vertebrates. (A**) Sequence alignment of Cdk8 (blue) and Cdk19 (green) proteins from: *Homo sapiens, Mus musculus, Xenopus tropicalis, Xenopus laevis, and Danio rerio*. Homologous sequences are black. The consensus (> 80%) is presented below the alignment. **(B)** Scheme indicating the fragment of *Cdk19* exon 1 removed by CRISPR-Cas9 (highlighted in red) in intestinal organoids. The arrow indicates the sequence of the sgRNA used. The sequence trace obtained after gene editing is presented below. The colour-code in the sequence in the box corresponds to the sequence trace.

**Fig S7. CDK8/CDK19 KO organoids are counter-selected**. WB indicating the levels of CDK8, CDK19 and phospho-Stat1-S727 in organoids after 14 days of tamoxifen treatment. Two out of the three *Cdk8* ^*-/-*^*/Cdk19* ^*-/-*^ clones (numbers 10 and 12) show a reappearance of the CDK8 protein: compare with Fig. 2B where proteins were extracted from the same samples, but one week earlier. (★) indicates the two clones where CDK8 protein is detected; this was observed only in organoids where double KO had been induced.

**Movie S1**. Live-cell microscopy shows a rapid expansion of both the lumen and total organoid surface area in WT organoids after the addition of forskolin. Cdk8^-/-^/Cdk19^-/-^ organoids do not swell after forskolin addition. Three different sizes of organoids are presented in each condition: big, (top); medium (middle) and small (bottom). Scale bar, 200 μm.

