## Supplementary figures and images for "CDK8 and CDK19 act redundantly to control the CFTR pathway in the intestinal epithelium"

### Supplementary figures

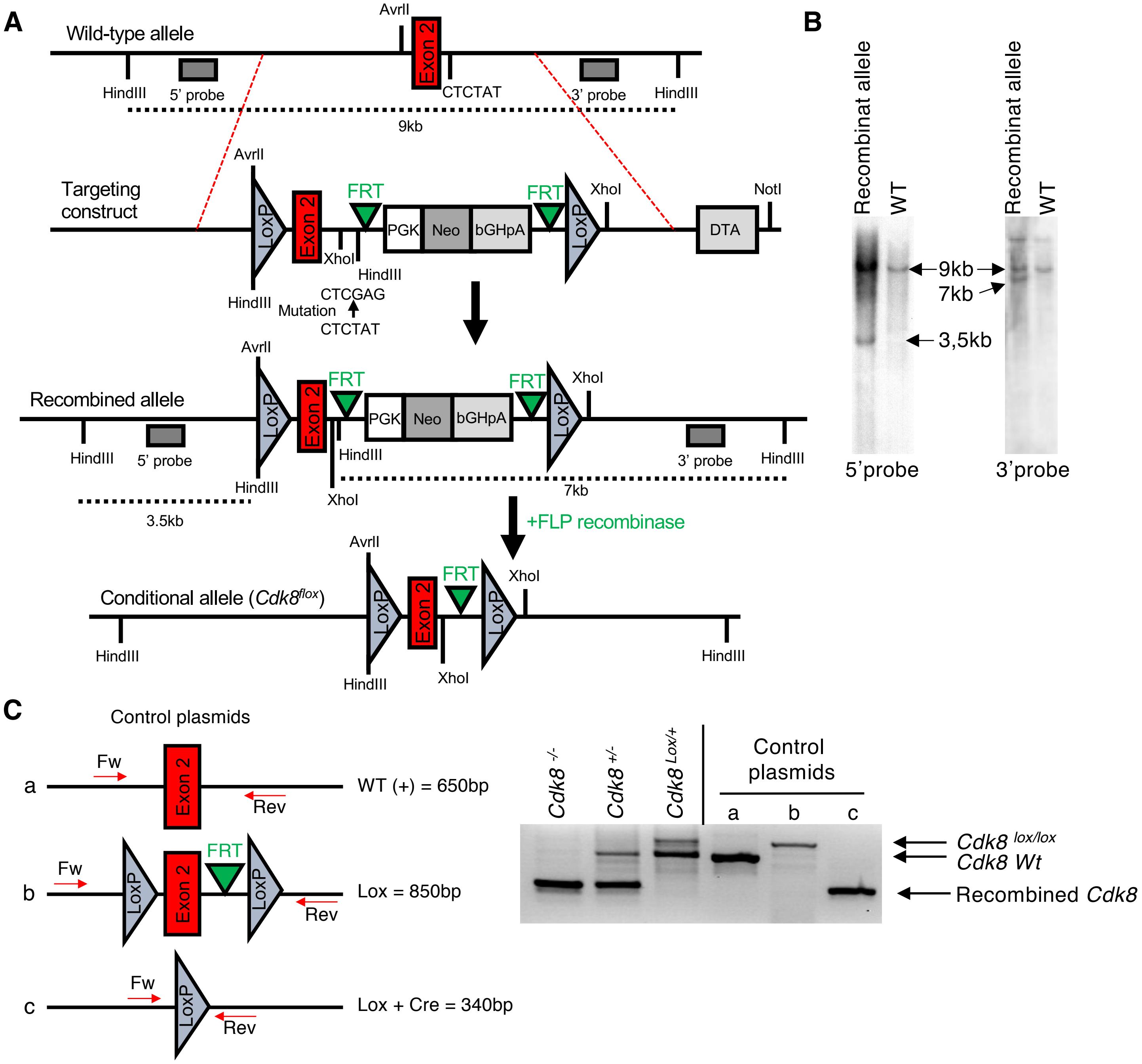

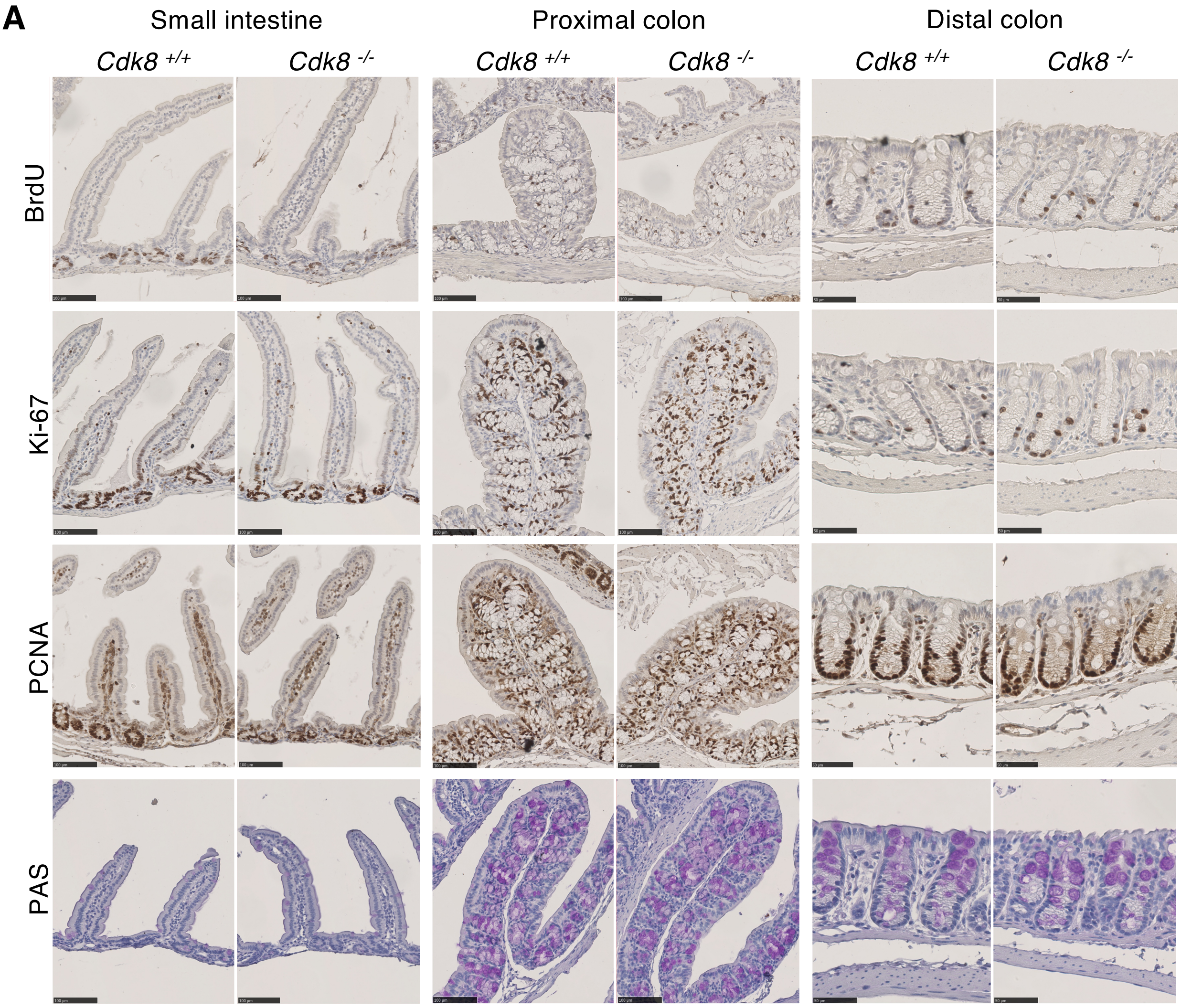

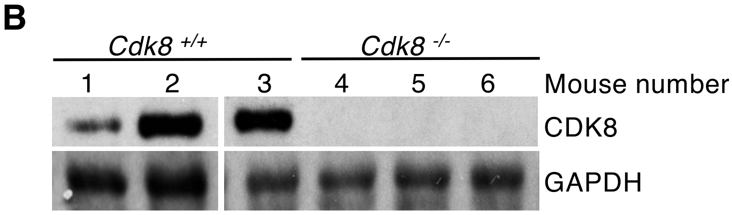

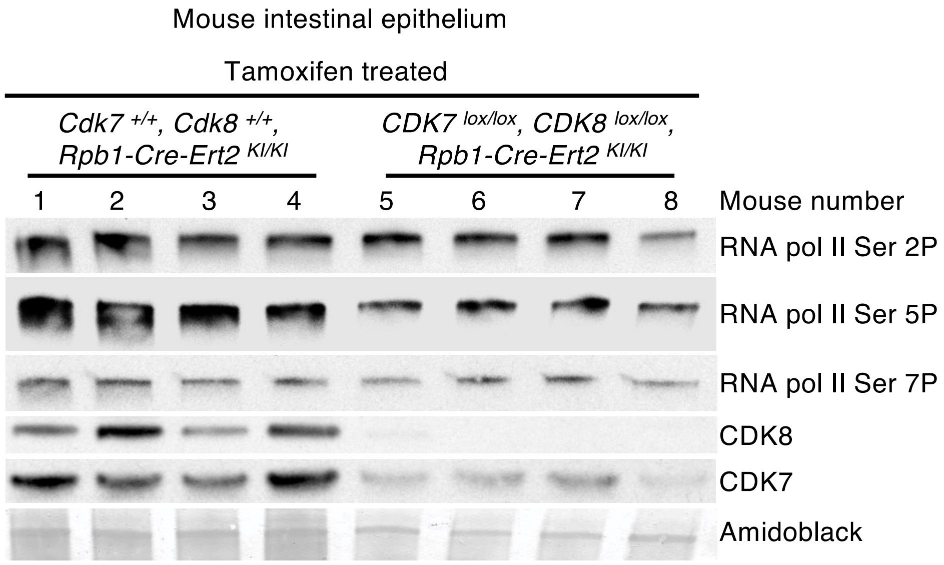

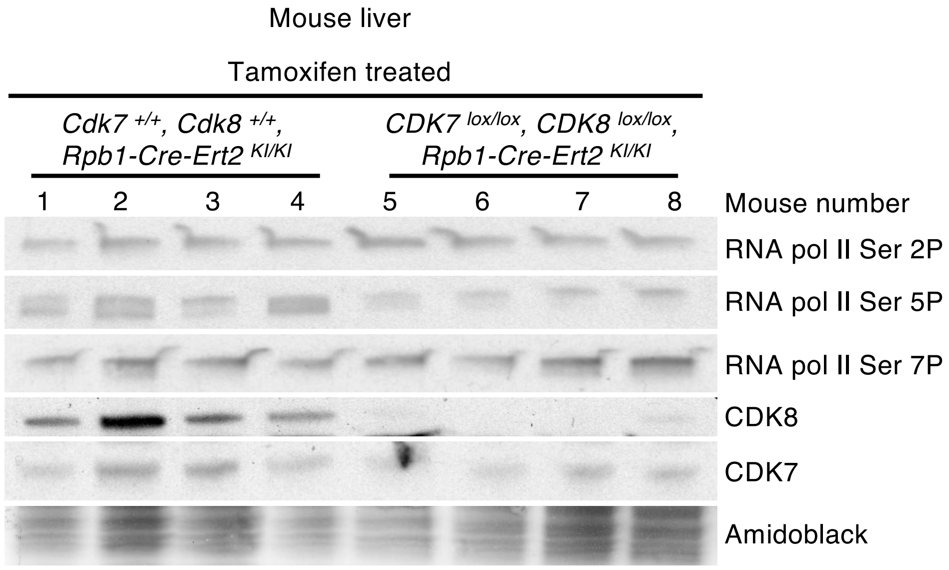

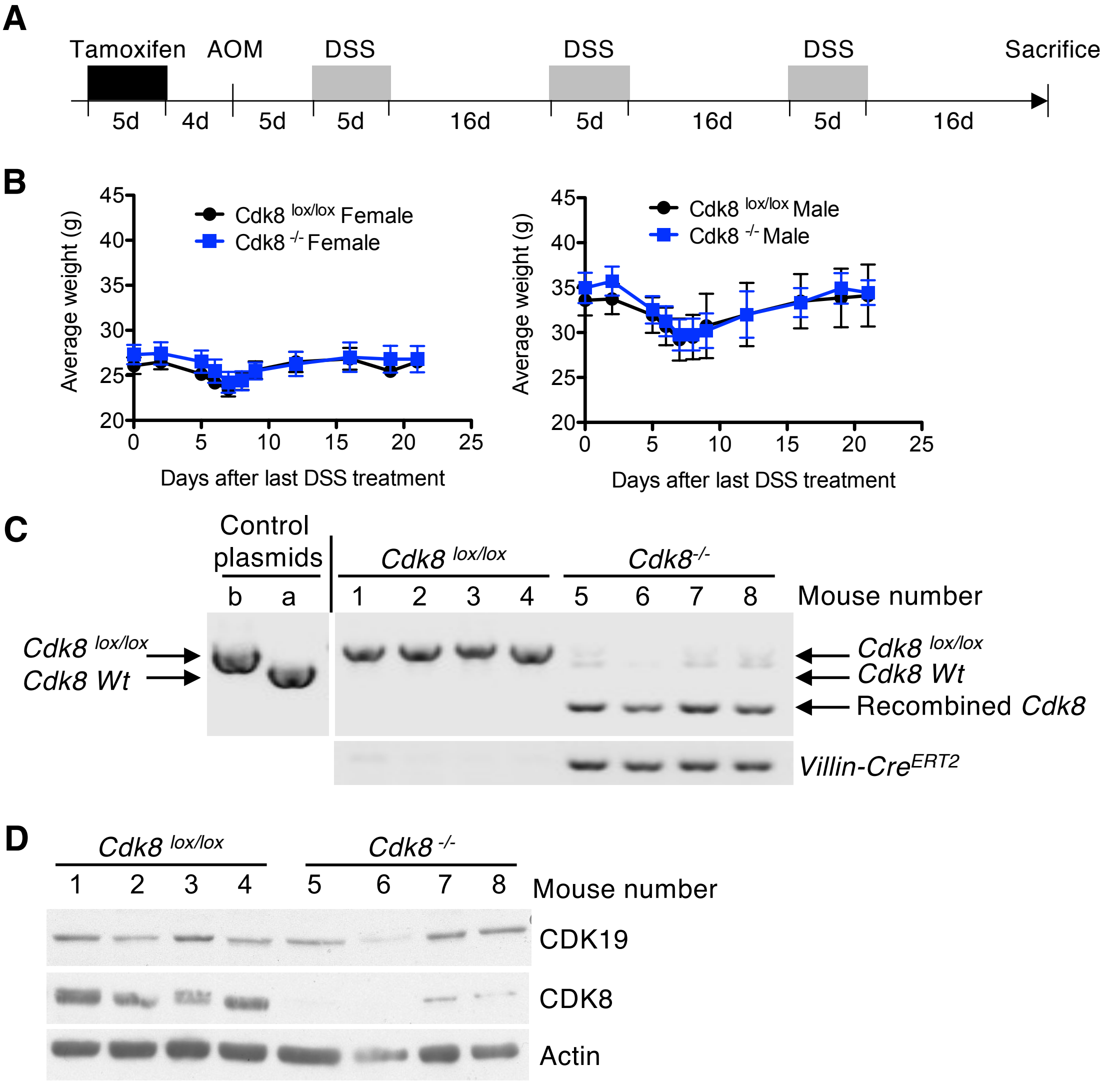

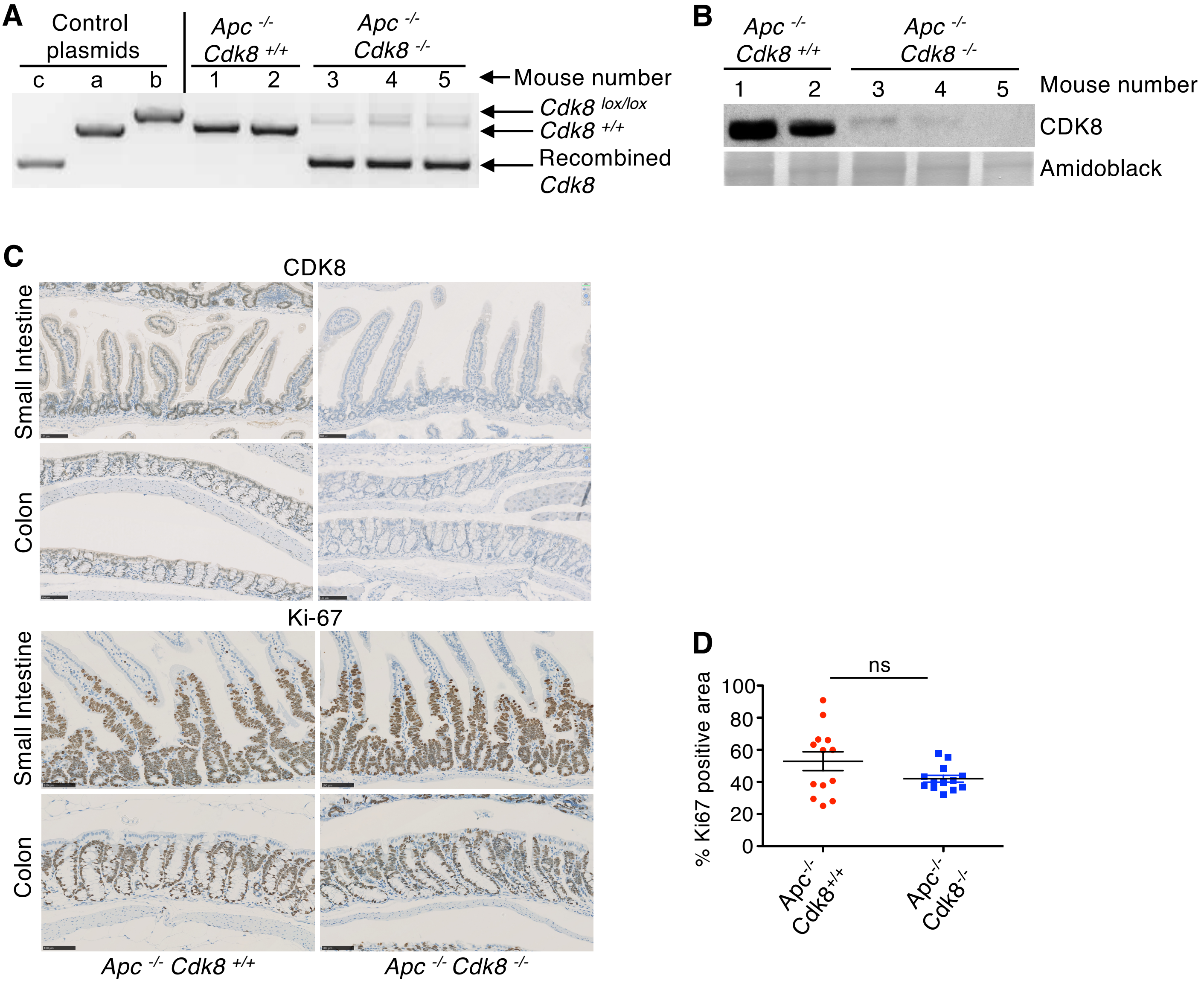

**Figure S6.**

**A**

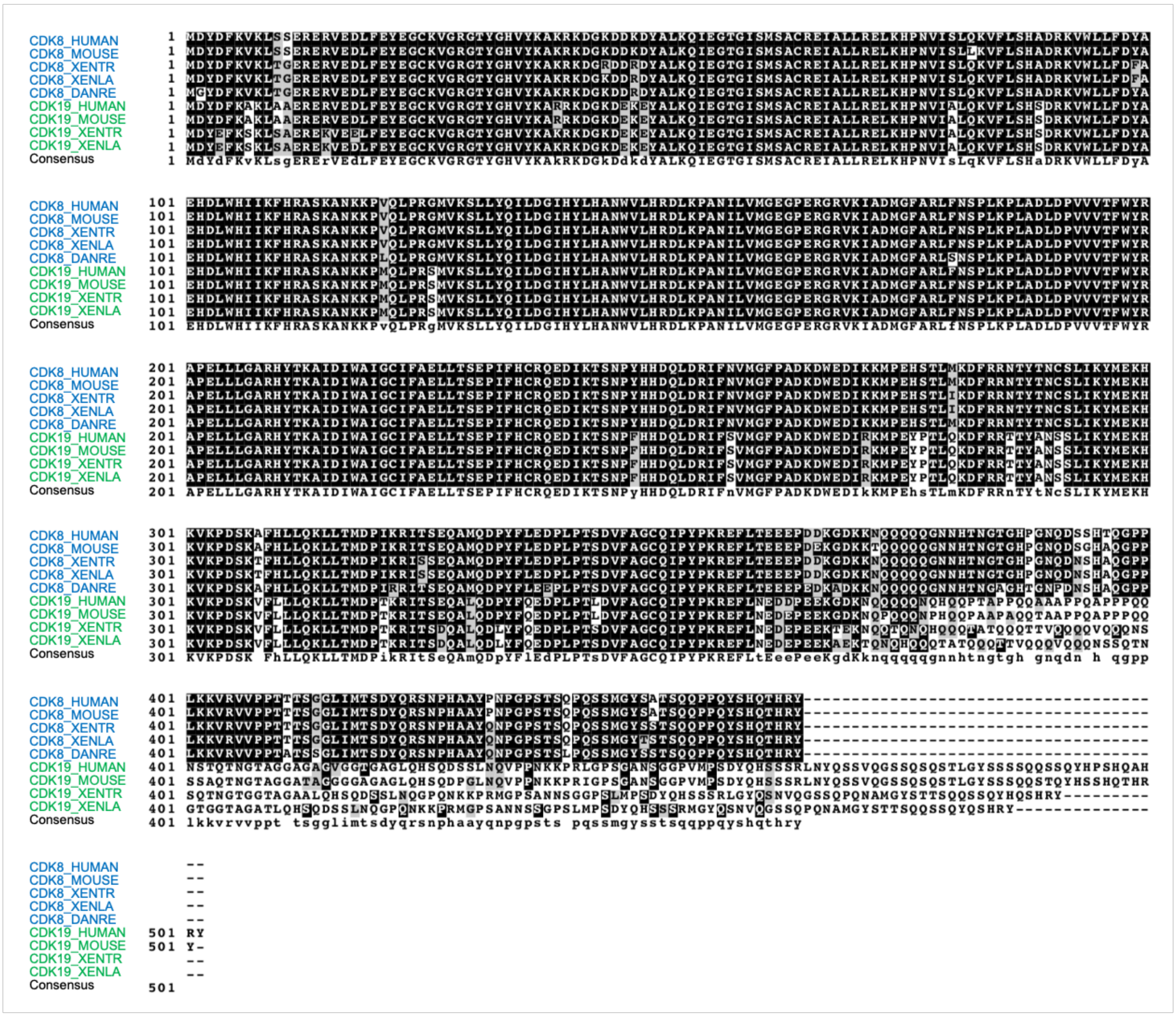


**B**

CRISPR-Cas9-targeted 40nt deletion in exon 1 of mouse *Cdk19*

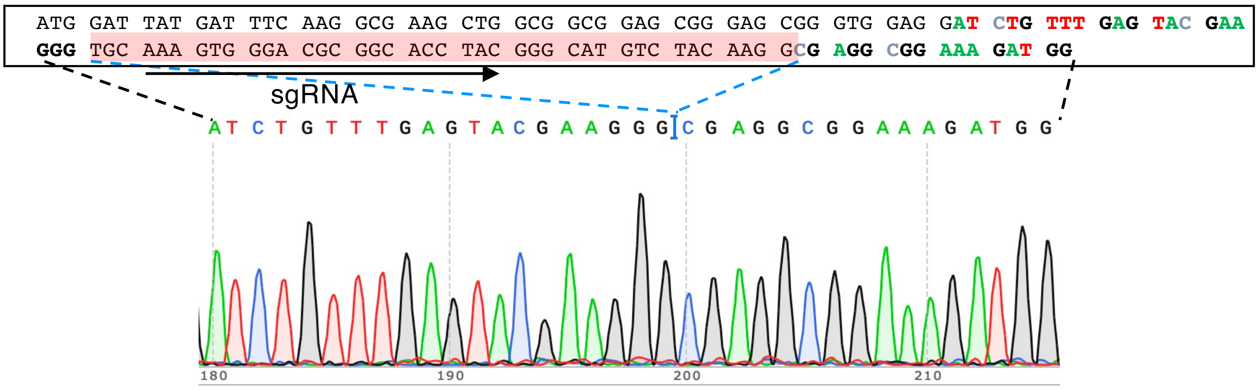

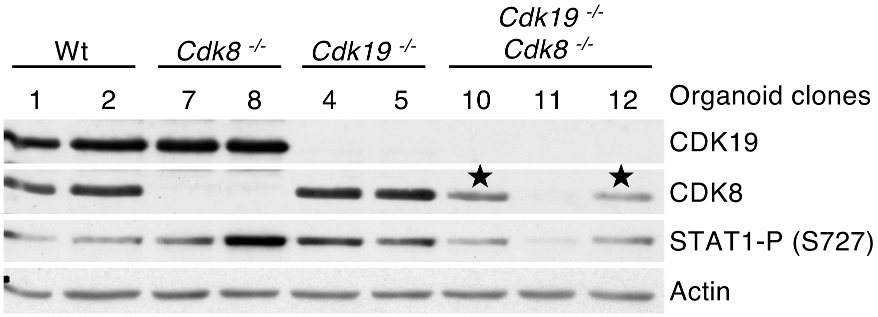
